# Tetramerization of Phosphoprotein is essential for Respiratory Syncytial virus budding while its N terminal region mediates direct interactions with the Matrix protein

**DOI:** 10.1101/2020.11.17.387951

**Authors:** Monika Bajorek, Marie Galloux, Charles-Adrien Richard, Or Szekely, Rina Rosenzweig, Christina Sizun, Jean-Francois Eleouet

## Abstract

It was shown previously that the Matrix (M), Phosphoprotein (P), and the Fusion (F) proteins of Respiratory syncytial virus (RSV) are sufficient to produce virus-like particles (VLPs) that resemble the RSV infection-induced virions. However, the exact mechanism and interactions among the three proteins are not known. This work examines the interaction between P and M during RSV assembly and budding. We show that M interacts with P in the absence of other viral proteins in cells using a Split Nano Luciferase assay. By using recombinant proteins, we demonstrate a direct interaction between M and P. By using Nuclear Magnetic Resonance (NMR) we identify three novel M interaction sites on P, namely site I in the α_N2_ region, site II in the 115-125 region, and the oligomerization domain (OD). We show that the OD, and likely the tetrameric structural organization of P, is required for virus-like filament formation and VLP release. Although sites I and II are not required for VLP formation, they appear to modulate P levels in RSV VLPs.

**Importance:** Human RSV is the commonest cause of infantile bronchiolitis in the developed world and of childhood deaths in resource-poor settings. It is a major unmet target for vaccines and anti-viral drugs. The lack of knowledge of RSV budding mechanism presents a continuing challenge for VLP production for vaccine purpose. We show that direct interaction between P and M modulates RSV VLP budding. This further emphasizes P as a central regulator of RSV life cycle, as an essential actor for transcription and replication early during infection and as a mediator for assembly and budding in the later stages for virus production.

## Introduction

Human RSV is the most frequent cause of infantile bronchiolitis and pneumonia worldwide (1). In France 460,000 infants are infected each year, of which ~30% develop lower respiratory infections and 4.8% to 6.7% are hospitalized, representing 45% of the young children admissions at the hospital (2). The enormous burden of RSV makes it a major unmet target for a vaccine and anti-viral drug therapy. However, despite over 60 years of research since its discovery, there is still no vaccine available, and RSV therapy remains mainly supportive. The current standard of care consists of prophylactic treatment of at-risk infants with Palivizumab (Synagis), a monoclonal antibody. However, its limited efficacy (approximately 50%), and high cost (€5.000 per treatment) limits its use to pre-term infants. As a result, 60% of at risk children remain untreated, and no efficient therapy is available to treat the adult population (3). There are currently 39 vaccines under development (4). One of the strategies for RSV vaccine development is based on virus like particles (VLPs). However, all anti-RSV VLP vaccines currently in preclinical development are using foreign viral systems incorporating the RSV glycoproteins (5, 6). This is mostly due to the inefficiency of RSV VLP production (most of the virus is cell-associated in cell culture (7)), and to insufficient understanding of RSV particle assembly and budding. RSV VLPs, if they can be produced at sufficient levels, will accurately mimic the viral morphology and structure. Identification of the minimal players involved in particle assembly and budding is an important step in understanding the mechanism behind RSV particle formation. The knowledge can be then used for large scale VLPs and attenuated virus production for vaccination purpose.

RSV belongs to the *Pneumoviridae* family in the order *Mononegavirales* (8). It primarily infects epithelial cells of the respiratory tract and replicates in the cytoplasm. It is an enveloped, non-segmented, negative-strand RNA virus. The viral genome is encapsidated by the nucleoprotein (N), forming a ribonucleoprotein (RNP) complex, which constitutes the template for the viral polymerase. It was recently shown that the replication and transcription steps of RSV take place in virus-induced cytoplasmic inclusions called inclusion bodies (IBs), where all the proteins of the polymerase complex, i.e. the viral polymerase (L), its main co-factor the P protein, the RNPs and the transcription factor M2-1 concentrate (9). It is noteworthy that pseudo-IBs, similar to those observed in RSV-infected cells, can be observed upon co-expression of only N and P (10, 11). We recently showed that the formation of these pseudo-IBs depends on a liquid-liquid phase separation induced by the N-P interaction (11). Once neo-synthesized, RNPs have to be exported from IBs to the plasma membrane, where RSV virions assemble and bud, forming elongated membrane filaments (12). According to the common paradigm, RSV assembles on the plasma membrane, and infectious viral particles are mainly filamentous (13, 14). However, recent data suggests that viral filaments are produced and loaded with genomic RNA prior to insertion into the plasma membrane. According to this model, vesicles with RSV glycoproteins recycle from the plasma membrane and merge with intracellular vesicles, called assembly granules, containing the RNPs (15, 16).

Regardless of the cellular location, the minimal RSV viral proteins required for efficient filament formation and budding of VLPs are P, M, and the F protein, more specifically its cytoplasmic tail (FCT) (17, 18). The atomic structure of the external part of F glycoprotein (excluding the transmembrane and cytoplasmic parts) has been resolved (19–21), but little is known about the FCT structure and its function in RSV assembly. M, a key structural protein, directs assembly and budding probably by interacting with FCT on the one hand, and with P associated to RNP on the other hand (22–24). M is required for filament elongation and maturation and, possibly, for transport of the RNP from IBs to the sites of budding (25). M was shown to localize to IBs where, presumably, the first interaction between M and the RNPs occurs. Some early reports have shown that M localization to IBs is mediated by interaction with M2-1 (26, 27). However, more recent work has demonstrated that M is found in IBs when expressed with the N and P proteins alone (17). As N is not required for RSV virus-like filament formation, M was suggested to interact with P. However, the exact mechanism of interactions between these proteins remains largely unknown. Structural data published previously by our group showed that M forms dimers and that the switch from M dimers to higher order oligomers triggers assembly of viral filaments and virus production (28). Based on M structure, a long patch of positively charged surface spanning the entire monomeric protein was suggested to drive the interaction with a negatively-charged membrane (29).

Functional and structural data are available for the P protein, which is a multifunctional protein capable of interacting with multiple partners. Recent studies allowed better characterization of its interactions and functions within the viral polymerase complex. P forms tetramers of elongated shape composed of a central oligomerization domain (OD), mapped to residues N131-T151 (30–34), and of N- and C-terminal intrinsically disordered regions (IDRs). Structural study of P in solution by NMR gave insight into the secondary structure propensity of these IDRs, forming almost stable helices in the C-terminal region and extremely transient helices in the N-terminal region (35). Residues 1-29 in the N-terminal region confer a chaperone function to P, by binding monomeric and RNA-free N (N^0^) and by maintaining N^0^ unassembled (36). P C-terminal residues 232-241 were shown to bind RNA-bound N assembled as rings mimicking the RNP (37). Very recently, the structure of the L protein bound to tetrameric P solved by cryo-electron microscopy revealed that each of the four P monomers adopts a distinct conformation upon binding to L, the entire L-binding region on P spanning residues 130-228. This includes the OD and the major part of the C-terminal domain (32, 38, 39). As part of viral transcription regulation, P region spanning residues 98-109 was shown to be the binding site for the RSV transcription anti-termination factor M2-1 (40). This interaction is involved in the recruitment of M2-1 to IBs (41). P also plays a pivotal role in M2-1 dephosphorylation mediated by the host protein PP1, which binds to P through an RVxF-like motif located at P residues 82-87 (41). Dephosphorylated M2-1 is recruited to specific regions in IBs, called IB associated granules (IBAGs), where viral mRNAs are concentrated, before trafficking back to the cytoplasm. This cycle is essential for RSV transcription and translation (41).

Recently, a P region encompassing residues 39–57 was found to be critical for VLP formation (42). In this study it was also observed that the OD of P had no significant contribution for VLP assembly and budding. Additionally, phosphomimetic substitutions in P region 39-57 inhibited VLP formation, suggesting that this region needs to be un-phosphorylated for VLP production (42). P region 110-120 was also shown to be required for efficient virus budding, affecting the final step of filament scission and virus release (42, 43). However, until now, no direct interaction between P and M has been shown. The lack of structural information on the putative M-P complex makes it difficult to speculate about the mechanism of budding. Thus, identifying specific RSV M-P protein-protein interactions has the potential to break new ground in our understanding of the mechanism behind RSV particle formation. This can further benefit RSV VLP-based vaccine research.

## Results

### Specific localization of M depends on expression of F, P, and N proteins

As shown previously, RSV VLPs can be generated independently of viral infection by transfecting cells with plasmids encoding the RSV M, P, N, and F proteins (18, 28). Although N is not required for RSV filament formation (17), it localizes with M in filaments when present, and is required, together with P, for the formation of pseudo-IBs (18, 28). In a first attempt to study the co-localization of M with N and P proteins in pseudo-IBs and in virus-like filaments at the plasma membrane, and to confirm the minimal requirement for filament formation in our system, we used a transfection-based assay.

BEAS-2B cells were transfected to express M, N, P, and F, or various combinations of these four proteins. The intracellular localization of RSV proteins and the formation of pseudo-IBs and virus-like filaments were determined by confocal imaging after staining in parallel with either anti-M and anti-N antibodies (Fig. 1A), or anti-P and anti-N (Fig. 1B). In the presence of M, N, P, and F, the formation of pseudo-IBs and of virus-like filaments was detected by immunostaining of M, N and P. Co-localization of M, N, and P was detected on filaments at the plasma membrane as well as in pseudo-IBs, as shown in zoomed merged images (Fig. 1A and 1B, row 1, positive control). When F was absent, M, P, and N localized in pseudo-IBs (Fig. 1A and 1B, row 2). M, but not P or N, was also found in small spikes, most probably at the plasma membrane. In the absence of P or N, no pseudo-IBs formed, as expected (Fig. 1A and 1B, rows 3 and 4). In the absence of P, M was found only as small spikes, whereas N spread all throughout the cytoplasm (Fig. 1A and 1B, row 3). Finally, as previously reported, large filaments containing M and P formed in the absence of N (Fig. 1A and 1B, row 4). Altogether, our results confirm previous observations showing that only M, P, and F are required for virus-like filaments formation, and that M co-localizes with N and P within pseudo-IBs and in virus-like filaments (17, 18, 26, 28).

**Fig.1:**
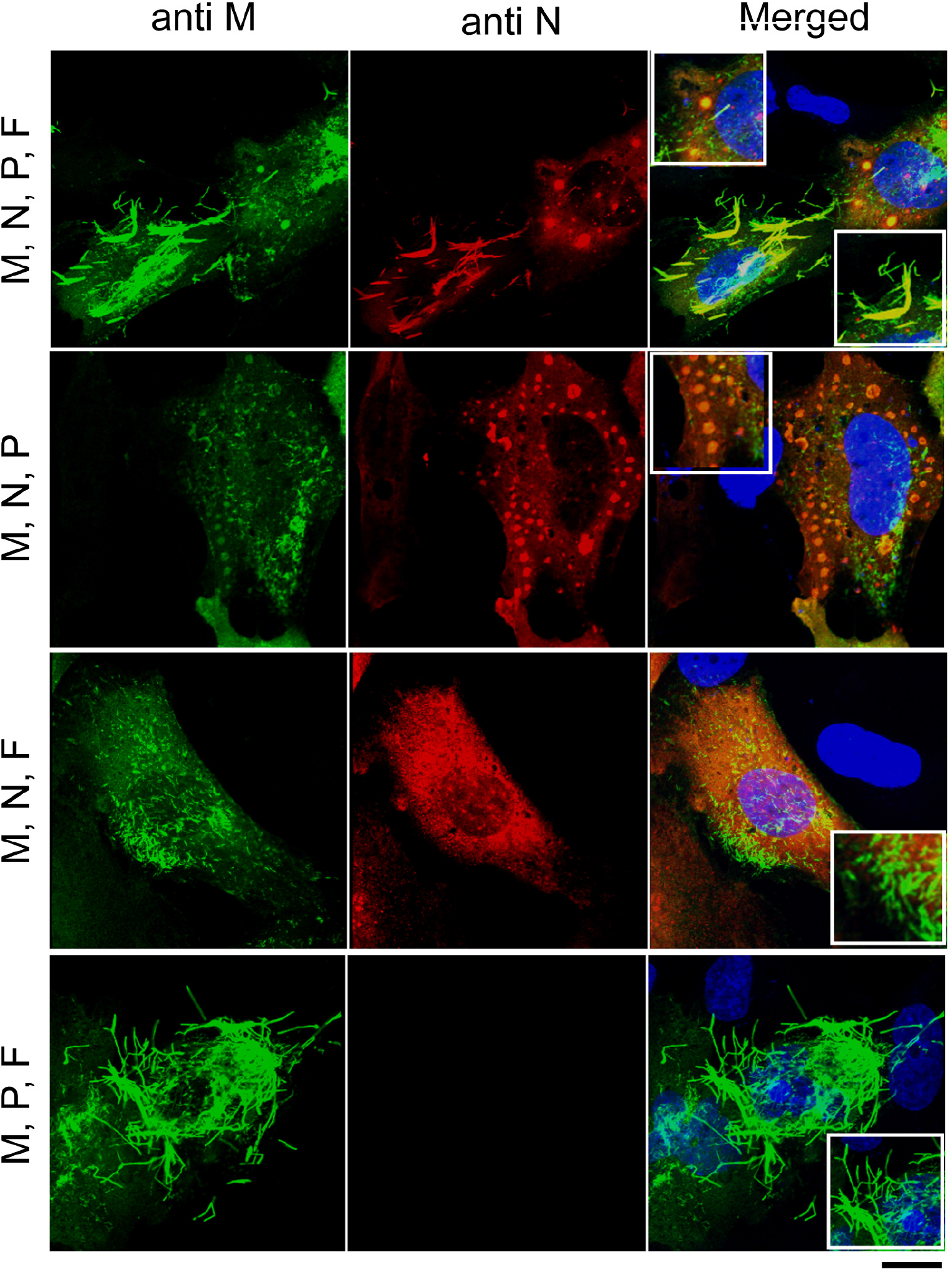
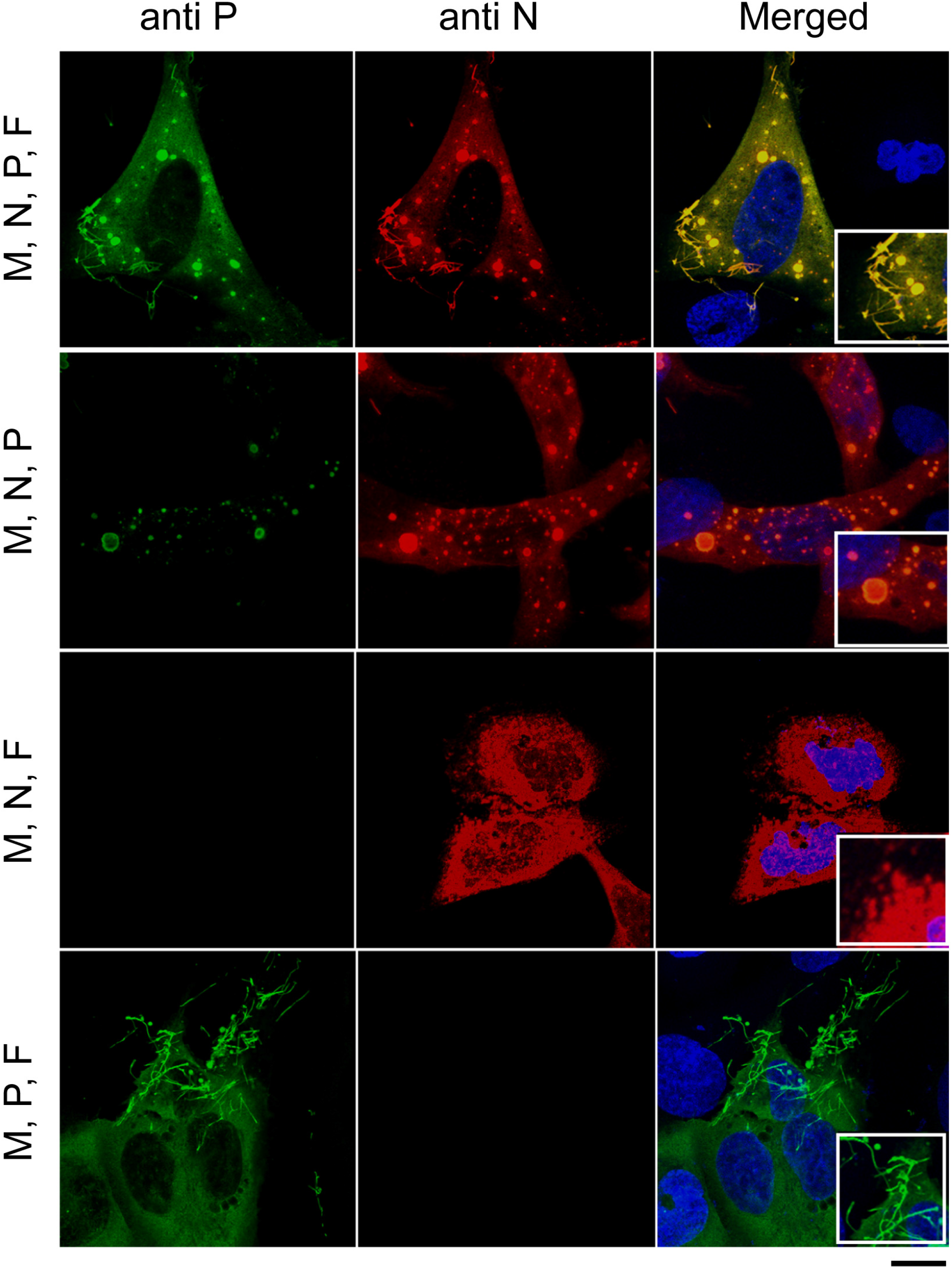
Specific localization of M depends on expression of F, P, and N proteins. BEAS-2B cells were co-transfected with pcDNA3.1 plasmids expressing RSV M, P, N, and F or a different combination of three proteins. 24 h post-transfection, cells were fixed, and immunostained with (**A)** anti-M (green) and anti-N (red) or (**B)** anti-P (green) and anti-N (red) primary antibodies followed by Alexa Fluor secondary antibodies, and were analyzed by confocal microscopy. Scale bars represent 10μm. Merged images are zoomed in 3X.

### M interacts with P in cells

As M localization in virus-like filaments seems to depend on P, we then studied whether M can interact with P in cells in the absence of other viral proteins. For that purpose, we used the NanoLUC interaction assay based on the split Nano Luciferase reporter (44). In this system, the 114 or the 11S Nano Luciferase fragments were fused to the C-terminus of viral proteins (Fig. 2A). Analysis of the lysates of transfected 293T cells by Western blotting using anti-P, anti-M, anti-N or anti-M2-1 antibody, confirmed protein expression with the fused NanoLUC fragments (Fig. 2B). In addition to the expected bands, a weak higher migrating band, probably unspecific, was detected in P114 and P-11S samples, whereas a double band was detected in M2-1-11S sample, corresponding most probably to the phosphorylated and unphosphorylated forms of M2-1 (41).To investigate the P-M interaction, combinations of two constructs were transfected into 293T cells. 24 h post transfection cells were lysed, luciferase substrate was added, and the luminescence, proportional to the strength of the interaction, was measured (Fig. 2C). As P is known to form tetramers (30–35), P/P interaction was used as positive control. As shown in Fig. 2C, transfection of P-114/P-11S resulted in a high luminescence signal, indicating a strong interaction. We also used the P-N interaction as a control: when co-expressing P-114 and N-11S, positive but relatively low luminescence was observed. Here, as the NanoLUC 114 subunit was cloned at the C-terminus of P protein, thus blocking the interaction between the C-terminus of P and RNA-bound N, the luminescence signal corresponds to the P-N^0^ interaction, which was previously shown to be rather weak, in the micromolar range (36). When M-114 was co-expressed with P-11S, the luminescence signal was similar to the P-N^0^ signal, suggesting that M interacts with P with similar affinity compared to P-N^0^ interaction. In contrast, expression of M-114 with M2-1-11S did not produce luminescence, suggesting that M and M2-1 did not interact when co-expressed in cells in our system. However, for this negative result, we cannot exclude that the C-terminal tag blocks protein-protein interactions that occur via the C-terminus of one or both proteins.

**Fig. 2:**
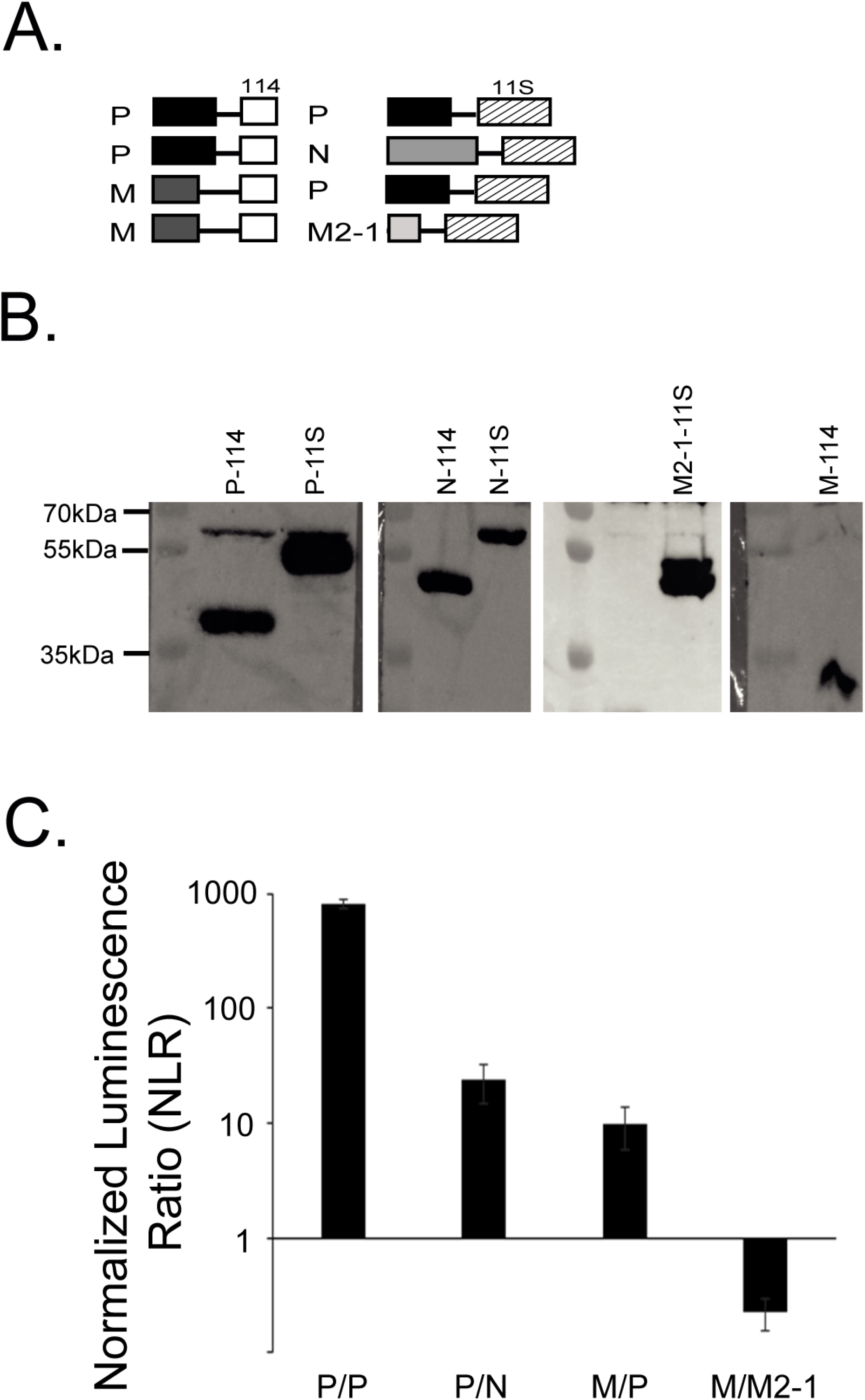
M interacts with P in cells. Protein-protein interactions were measured using the NanoLuc assay. **(A)** Scheme of the RSV protein constructs fused with NanoLUC 114 or 11S subunit and pair combinations used in (C). **(B)** 293T cells were transfected with plasmids encoding P, N, M, or M2-1 fused to 114 or 11S NanoLUC subunit. Cells were lysed 24 h post transfection, and cell lysates were then subjected to Western analysis using anti-P, anti-M, anti-N or anti-M2-1 polyclonal antibody. Size markers are shown on the left side of each gel. **(C)** 293T cells were transfected with pairs of constructs, combined as shown in the graph. P/P and P/N were used as positive controls. Cells were lysed 24 h post transfection, and luminescence was measured using a Tecan Infinite 200 plate reader. The NLR is the ratio between actual read and negative controls (each protein with the empty NanoLUC vector). The graph is representative of four independent experiments, each done in three technical repeats. Data represents the means and error bars represent standard deviation across 4 independent biological replicates.

### M directly interacts with P via its N-terminal region and OD

Next, we investigated whether M directly interacts with P *in vitro* using recombinant proteins, and determined the P region involved in the interaction. Based on structural data available for isolated P (30, 35), we generated rational deletions of each subdomain of P (Fig. 3A). While P fragments lacking the OD (P_ΔOD_, P_1-126_ and P_161-241_) were shown to be monomeric by NMR, P fragments containing the OD (full-length P, P_OD_, P_1-163_ and P_127-241_) appeared to be associated via the OD (35). The purity and the size of all the P fragments were controlled by SDS-PAGE (Fig. 3B). M and full-length P or P fragments were co-incubated and the formation of complexes was analysed by native agarose gel electrophoresis. Of note, in native gels proteins migrate according to the combination of their size, shape and charges. This explain why P_ΔOD_ migrates at higher apparent molecular weight than full-length P, as the global charges are −20.7 for monomeric P_ΔOD_ and −86.8 for tetrameric P at pH 7.4. When M was incubated with P fragments, shifts were observed only with full-length P and P_1-163_ (Fig. 3C), indicating that the N-terminal domain of P (residues 1-131) and the OD (residues 131-151) are together required for a stable P-M interaction in this system. Additionally, no shift was observed neither for P_ΔOD_ nor for P_1-126_ when incubated in the presence of M. This suggests that either the M-binding site is at least partly on the OD of P, or that M interaction requires tetrameric P. Since no shift was observed for P_OD_, P_127-241_ and P_161-241_ when incubated with M, these results showed that neither the OD nor the C-terminal region of P on their own or together were sufficient for a detectable P-M interaction *in vitro.*

**Fig. 3:**
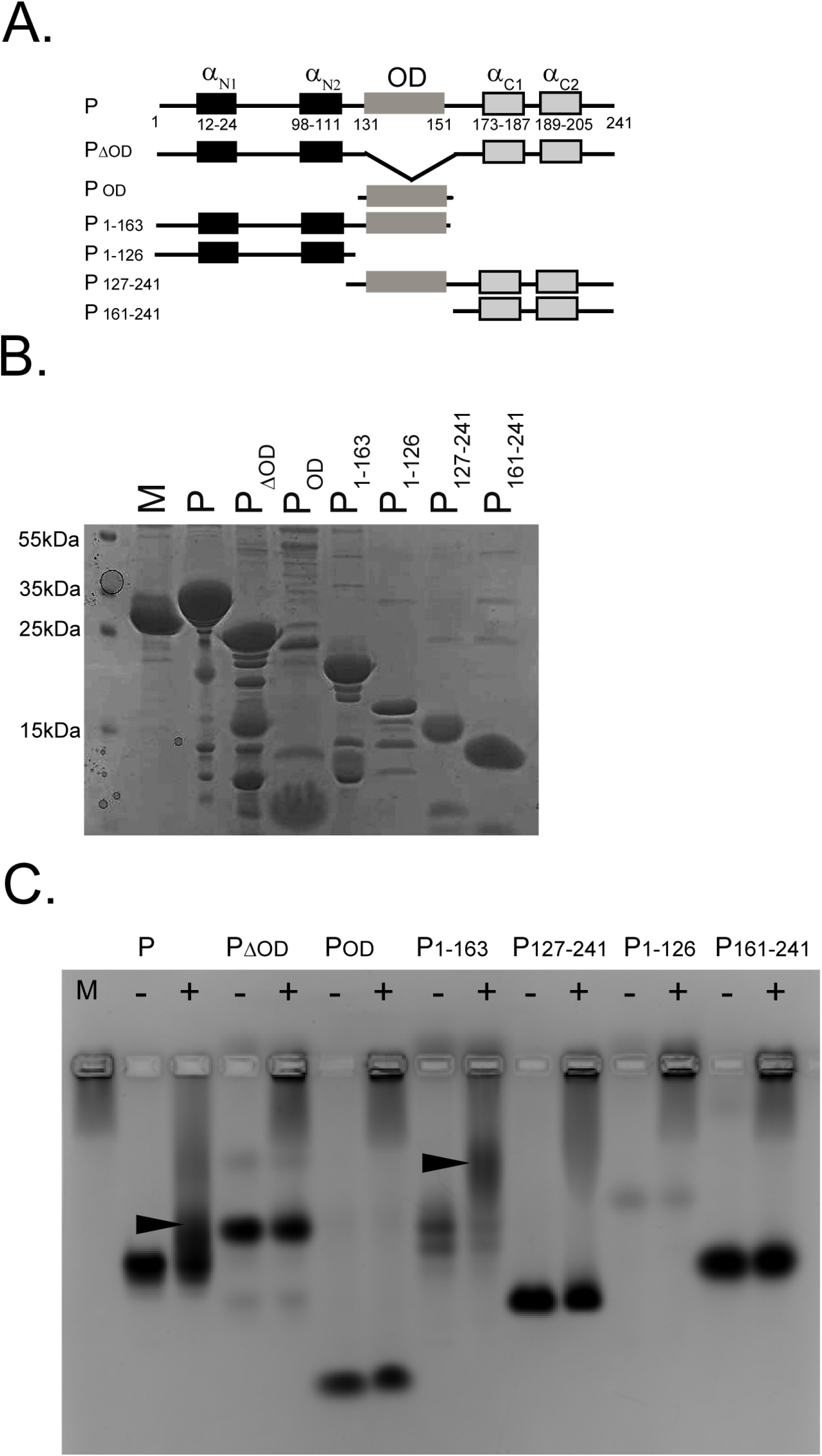
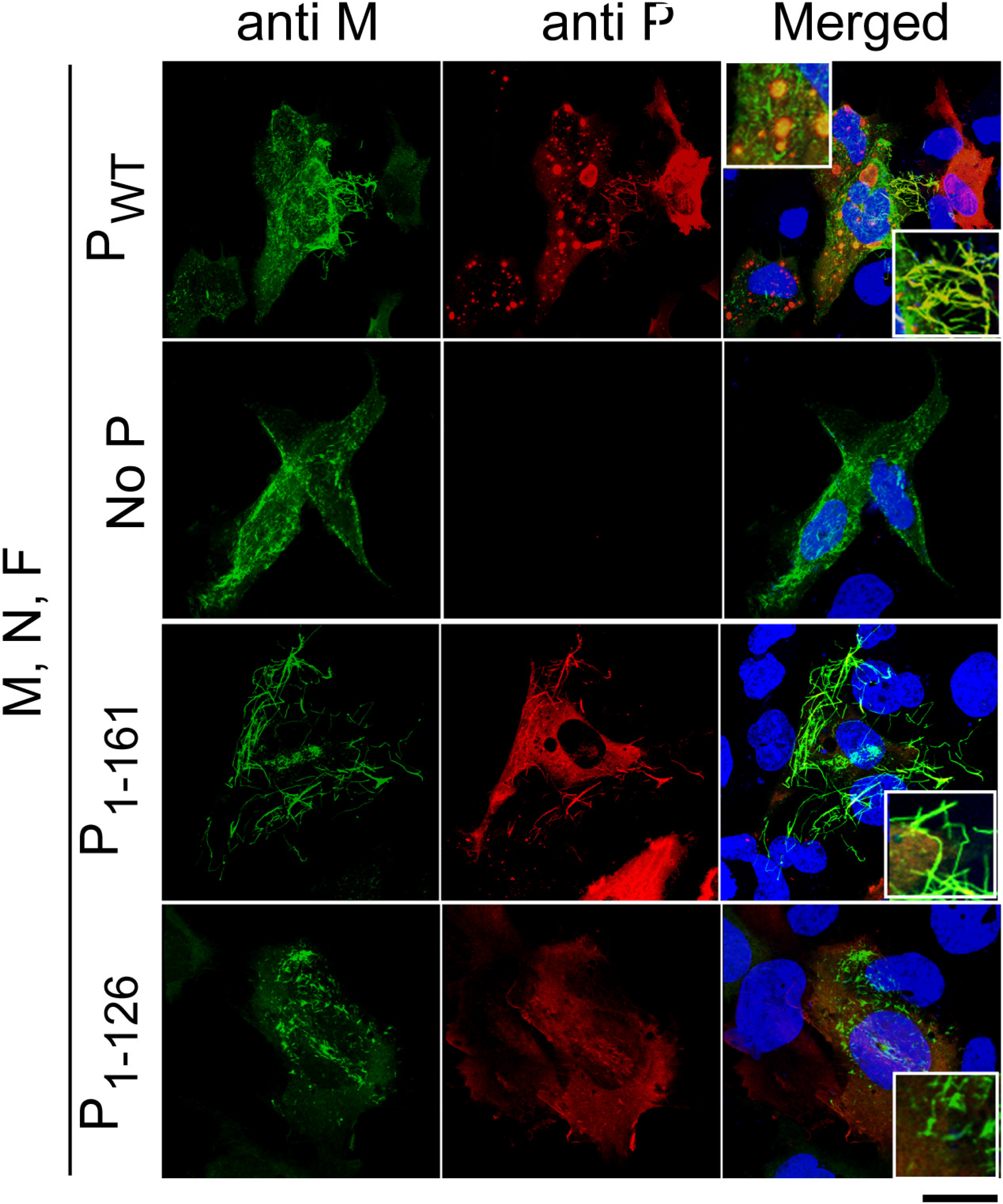
M directly interacts with P via its N-terminal region and OD. **A.** Scheme of the secondary structure of P protein (α_N1_, α_N2_, α_C1_ and α_C2_ denote transient helices detected in isolated P (35)) and the P constructs used. P_1-163_ and P_1-161_ constructs were used for bacterial and mammalian expression, respectively. **B.** SDS-PAGE and Coomassie blue staining of purified recombinant M, P and P fragments. **C.** M protein was co-incubated with P or P fragments for 30 min prior analysis of formation of complexes by band shift on native agarose gel. Arrows indicate complex formation. **D.** BEAS-2B cells were co-transfected with pcDNA3.1 plasmids expressing RSV M, F, N, and P WT or P deletion mutants. Cells were fixed, and immunostained with anti-M (green) and anti-P (red) antibodies followed by Alexa Fluor secondary antibodies, and were analyzed by confocal microscopy. Scale bars represent 10μm. Merged images are zoomed in 3X.

Based on these results, we next wanted to assess if a P fragment containing the N-terminal region and the OD was sufficient to interact with M and to induce membrane filaments in cells. We used the P_1-161_ fragment, shorter than P_1-163_ by two residues, which are outside of the OD on the C-terminal side. Again, we used the filament formation assay. BEAS-2B cells were transfected to express M, N, F and P, P_1-161_ or P_1-126_ constructs, and the formation of RSV virus-like filaments was determined by confocal imaging after staining with anti-M and anti-P primary antibodies (Fig. 3D). As previously shown (Fig. 1), in the presence of M, N, F, and P, the formation of pseudo-IBs and virus-like filaments was observed, and M co-localized with P in both structures (Fig. 3D, row 1, positive control). Co-localization was not seen in all pseudo-IBs, but this could reflect different maturation states of pseudo-IBs. In the absence of P or when P_1-126_ was expressed, neither IBs nor virus-like filaments were detected, and M was found in small spikes (Fig 3D, row 2 and 4). Virus-like filaments were detected when the P_1-161_ construct was expressed, and M-P_1-161_ co-localization occurred in virus-like filaments (Fig 3D, row 3). These results are in agreement with those obtained with the gel shift assay using recombinant proteins (Fig. 3C), and confirmed that the P_1-161_ fragment is competent for M binding and for filament formation. It is noteworthy that under these conditions no pseudo-IBs were detected, because the interaction between the C-terminus of P (missing in the P_1-161_ construct) and N is critical for their formation (45). Altogether, our results show that the P_1-161_ fragment is sufficient to interact with M and to induce the formation of virus-like filaments in cells in presence of M, N, and F.

### Identification of M interaction sites on P by NMR

Next, we sought for a method to identify the M interaction site on P, in a residue– specific manner, without resorting to mutations or internal deletions in potential regions of interest. We have previously used NMR to localize molecular recognition features (MoRFs) on P, taking advantage of the intrinsically disordered nature of this protein (46). Perturbations in amide ^1^H-^15^N correlation spectra, either of intensities (Fig. 4A and 5) or chemical shifts, were used to map regions that sense direct interaction and/or conformational changes due to binding of a partner.

**Fig. 4:**
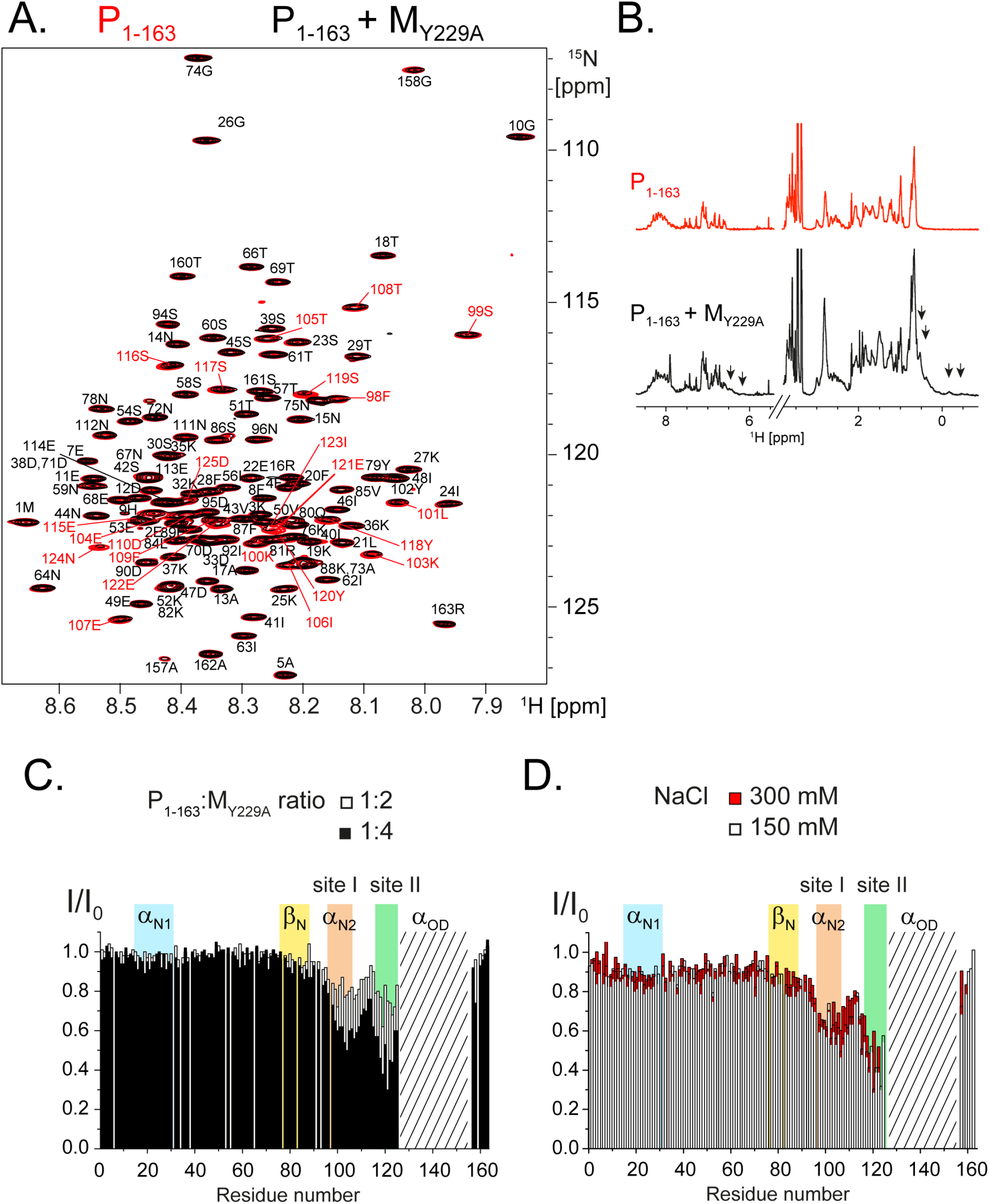
Observation of a direct interaction between RSV P1-163 and M-Y229A mutant by NMR. **A.** Superimposed 2D ^1^H-^15^N BEST-TROSY HSQC spectra and **B.** 1D ^1^H spectra with water suppression of 25 μM ^15^N-labeled P_1-163_ alone (red) and with 4 molar equivalents of M-Y229A (black). Acquisition was done in 50 mM Na phosphate pH 6.7, 150 mM NaCl buffer at 800 MHz ^1^H frequency and a temperature of 288 K. Residue-specific assignment of each 2D peak is indicated by the P residue number and the amino acid type. Signals with significant intensity decrease are annotated in colour. Arrows indicate specific M ^1^H NMR signals, in methyl (-1-1 ppm) and aromatic (6-7 ppm) proton regions. **C**. Intensity ratios (I/I_0_), represented as bar diagrams, were measured for each peak in the HSQC spectra of 25 μM ^15^N-labeled P_1-163_ in the absence and in presence of M-Y229A, at P:M molar ratios of 1:2 and 1:4. Signals from the α-helical oligomerization domain (α_OD_, hatched area) are broadened beyond detection. Coloured background indicates the localization of P specific regions: transient helices α_N1_, α_N2_ (also termed site I), extended region β_N_ and site II upstream of α_OD_. **D.** Intensity ratios (I/I_0_) measured from HSQC spectra of 40 μM P_1-163_ in the absence and in the presence of 150 μM M-Y229A, using two different salt concentrations, 150 mM and 300 mM, as indicated.

**Fig. 5:**
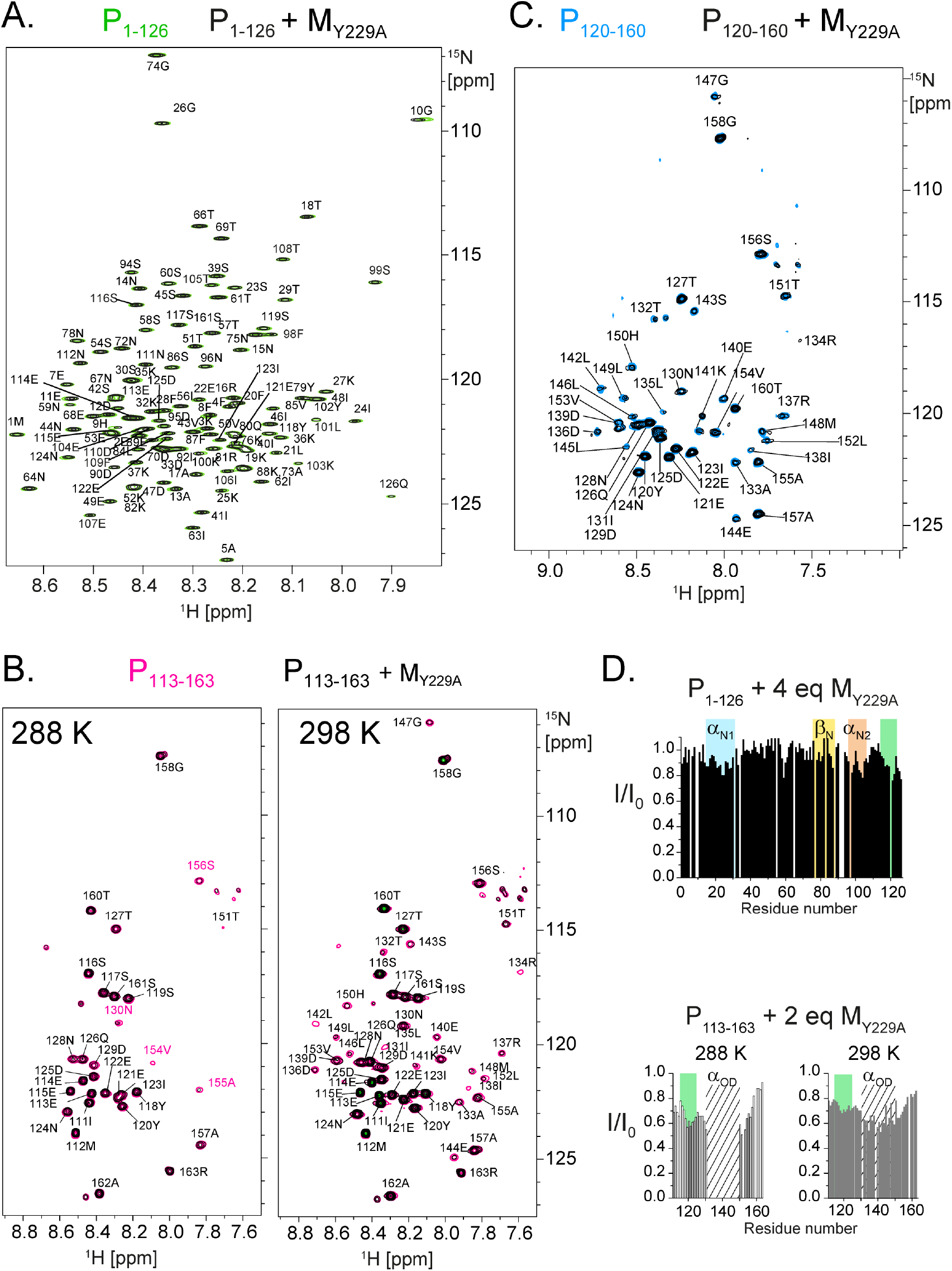
Localisation of RSV M-Y229A interaction regions on P by NMR using fragments of P_1-163_. Superimposed 2D ^1^H-^15^N BEST-TROSY HSQC spectra of ^15^N-labeled N-terminal P fragments. Samples were in 50 mM Na phosphate pH 6.7 150 mM NaCl buffer. Spectra were recorded at 800 MHz ^1^H frequency. **A.** 25 μM ^15^N-labeled P_1-126_ alone (green) and after addition of 4 molar equivalents of M-Y229A (black) at a temperature of 288 K, **B.** 50 μM ^15^N-labeled P_113-163_ alone (pink) and with 2 molar equivalents of M-Y229A (black), at 288 K and 298 K, **C.** 25 μM ^15^N-labeled P_120-160_ alone (blue) and in the presence of 2 molar equivalents of M-Y229A (black), at 288 K. **D.** Intensity ratios (I/I_0_) were determined from the HSQC spectra of P_1-126_ and P_113-163_ with and without M-Y229A. Signals from the OD (α_OD_, hatched area), which are broadened beyond detection at 288K for P_113-163_, become visible at 298 K. Other areas are highlighted using the same colour code as in Fig. 4.

To control the oligomerization state of M, we used the M-Y229A mutant, which was shown to form dimers, but is less prone to form higher order oligomers as compared to WT M (28). No aggregation or self-assembly of M-Y229A was observed, as the samples stayed clear and M-specific signal was detected in ^1^H NMR spectra, also in the presence of P (Fig. 4B). Of note, since M-Y229A is a dimeric 2×29 kDa folded protein, M signals are much broader than P signals that stem from the intrinsically disordered N-terminal region of P. This was also verified by SEC analysis made after NMR measurements (data not shown). A buffer with reduced ionic strength (150 mM NaCl as compared to 300 mM, the usual M storage buffer (13)) was used to enhance potential electrostatic interactions between M and P. The temperature was set to 288 K to increase the stability of M in this buffer, and also to reduce NMR signal broadening of solvent-exposed amide protons due to exchange with water. The final concentration of M was set to 100 μM. Concentrations and molar ratios are given for protomers, independently of the oligomerization state of the proteins. Since the C-terminal part of P seemed to be dispensable for interaction with M and for virus-like filament formation (Fig. 3), we performed experiments with ^15^N-labeled P fragments devoid of this part rather than with full-length P: this reduces signal overlap in ^1^H-^15^N correlation spectra.

Incubation of ^15^N-labeled P_1-163_ with M-Y229A at a P:M molar ratio of 1:4 (25 μM P and 100 μM M) resulted in significant intensity decrease of several P_1-163_ amide signals in 2D ^1^H-^15^N correlation NMR spectra (Fig. 4A). These perturbations took place in two proximal regions located immediately upstream of the OD: in the α_N2_ region (residues 98-111) and in a stretch spanning residues 115-125 (Fig. 4C). The first corresponds to the extremely transient helix α_N2_, previously identified as the binding site of RSV M2-1 protein (46, 47). Due to specific dynamics in the α-helical coiled-coil OD, amide ^1^H-^15^N signals are broadened out beyond detection at 288 K for P fragments that contain the OD flanked by N- and/or C-terminal extensions (46). Hence, M-binding to the OD could not be assessed using P_1-163_.

A concentration effect for P_1-163_ was evidenced by comparing the intensity ration I/I_0_ measured with two different P:M molar ratios (Fig. 4C). A maximal effect, where signals disappear completely, could not be reached, because M could not be concentrated above 100 μM without starting to aggregate and because the concentration of ^15^N-labeled P had to be kept >10 μM due to the sensitivity limitation of NMR. Altogether, the NMR experiments with P_1-163_ show that an interaction takes place between P_1-163_ and M-Y229A, which involves a region immediately upstream of the OD. This interaction is rather weak. A set of NMR data with P_1-163_ and M-Y229A was also acquired in a buffer with 300 mM NaCl and compared to 150 mM NaCl (Fig. 4D). Overall intensity perturbation is larger with lower salt concentration, indicating that M-Y229A binding is stronger under these conditions and suggesting that there may be an electrostatic contribution to the P-M interaction.

To assess the role of the OD, we used the P_1-126_ fragment, which is devoid of it. When incubated with 4 molar equivalents of M-Y229A, no significant effect was observed in the intensities of the ^1^H-^15^N correlation spectrum (Fig. 5A and 5D, upper panel), indicating that the OD is required for M binding. These results correlate well with the band shift experiments for the M/P complex using native gel analysis (Fig. 3C). To further characterize the binding site on P_1-163_, we performed NMR interactions experiments using the shorter P_113-163_ fragment, which contains only the OD flanked by residues 115-130 and 152-163 (Fig. 5B and 5D, lower panel). In the presence of M-Y229A, NMR signal intensities decreased by 30 % upstream and downstream of the OD. A stronger effect was again observed for residues 115-125 (Fig. 5B and 5D, lower panel), suggesting that they play a particular role in M binding. Like for P_1-163_, the OD could not be observed for P_113-163_ fragment at 288 K. However, these signals become visible at higher temperature for P_113-163_ (Fig. 5B and 5D, lower panel). The intensity perturbation pattern of P_113-163_ at 298 K suggests that the OD region is affected by the presence of M-Y229A. For P_120-160_, the smallest fragment used in this study, all amide signals can be observed, even at 288 K (Fig. 5C). M-Y229A induces a weak global reduction in intensity that may point to weak binding to P_120-160_. Taken together, these results suggest that M binding to P is achieved through multiple contact sites that are located in α_N2_ (site I), in the 115-125 region (site II) and potentially in the OD. Binding to any of these sites appears be rather weak, and M binding to the two N-terminal P regions requires tetrameric P.

### Validation of M-binding sites on P using P deletion mutants

In order to validate the NMR data showing multiple M binding sites, we then performed band shift assays with different fragments of P. We generated P deletions using P_1-163_ as a template and deleted a different part of the N-terminal domain for each construct. The OD was present in all constructs to keep the P protein as a tetramer (Fig. 6A). Each construct contained different M-binding domains. P_40-163_ contained the region spanning residues 39-57, previously reported to affect RSV assembly and potentially being involved in M interaction (42), as well as all binding sites identified by NMR (Fig. 4 and 5). P_80-163_ contained the two potential binding sites I and II identified by NMR. P_113-163_ contained only site II, and P_120-160_ lacked both sites, containing only the OD. We used both WT M and M-Y229A in order to ascertain that the NMR results were not biased by the Y229A M mutation. M proteins, full-length P and P fragments were purified as previously and analysed by SDS-PAGE (Fig. 6B). M and P constructs were co-incubated before analysis of complexes by native agarose gel electrophoresis (Fig. 6C). Shifts were observed with full-length P and P_1-163_, as shown above in Fig. 3C. P_40-163_ and P_80-163_ fragments (containing both sites I and II and OD) also induced a shift. No band shift was detected when M was incubated with P_113-163_ (containing only site II and OD) or with P_120-160_. Similar shifts were observed for M WT and M Y229A. This is in line with our NMR results (Fig. 4 and 5), indicating that M binding by P is achieved through multiple sites. Interactions seen by NMR with P_113-126_ or P_120-160_ were not strong enough to induce a band shift when analyzed by native gel electrophoresis.

**Fig. 6:**
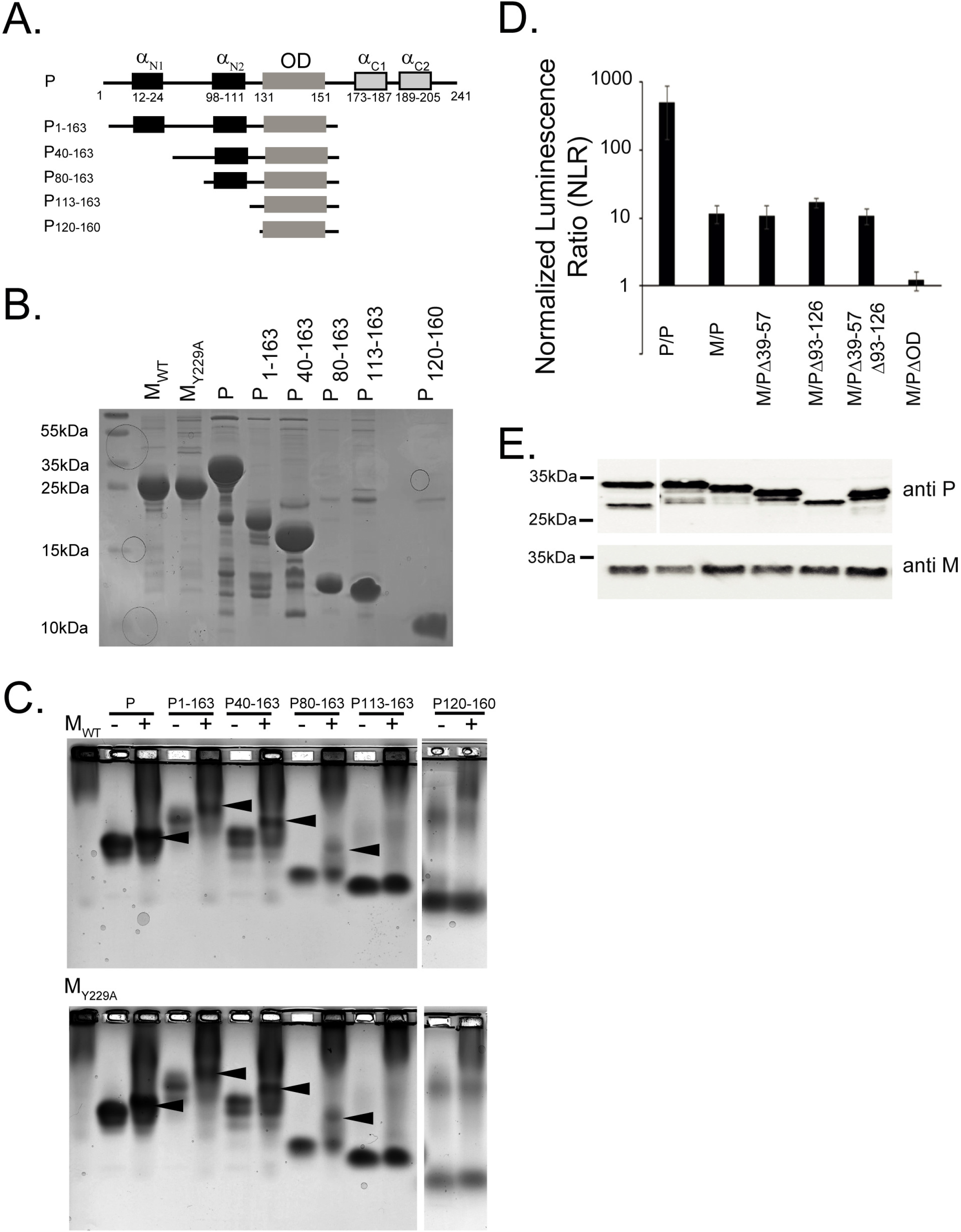
Validation of novel M-binding sites on P using P deletion mutants. **A.** Scheme of the P protein secondary structure and the P constructs used. **B.** His-tagged and size-exclusion chromatography purified M, P or P fragments purified on glutathione beads were analysed using SDS-PAGE and Coomassie Blue staining. **C.** Purified M WT (upper panel) or M Y229A (lower panel), and P or P deletion constructs were incubated 30 min prior analysis of formation of complexes by band shift on native agarose gel. Arrows indicate complex formation. **D.** 293T cells were transfected with pairs of M and P constructs fused to 114 or 11S NanoLUC subunit, combined as shown in the graph. P/P was used as positive control. Cells were lysed 24 h post transfection, and luminescence was measured using a Tecan Infinite 200 plate reader. The NLR is the ratio between actual read and negative controls (each protein with the empty NanoLUC vector). The graph is representative of four independent experiments, each done in three technical repeats. Data represents the means and error bars represent standard deviation across 4 independent biological replicates. **E.** Same cellular lysates used in D. were then subjected to Western analysis using anti-P or anti-M polyclonal antibody. Size markers are shown on the left side of each gel.

In order to verify which of the identified specific M/P binding regions is relevant for interaction between M and P in cells, we used again the NanoLUC interaction assay based on the split Nano Luciferase reporter. The 114 or the 11S Nano Luciferase fragments were fused to the C-terminus of M and P proteins (Fig. 2A). P_WT_ and four P deletion mutants were analysed: P_Δ39-57_ (deleted of the region previously reported to be involved in M/P interaction (42)), P_Δ93-126_ (lacking sites I and II as identified by NMR), P_Δ39-57_,_Δ93-126_ (lacking the three regions), and P_ΔOD_. Combinations of two proteins were transfected into 293T cells. 24 h post transfection cells were lysed, luciferase substrate was added, and the luminescence, proportional to the strength of the interaction, was measured (Fig. 6D). P/P interaction was again used as positive control. As shown in Fig. 6D, transfection of P-114/P-11S resulted in high luminescence signal, indicating a strong interaction. Transfection of M-114 and P-11S resulted in positive signal (comparable to Fig. 2C). Transfection of P_Δ39-57_-11S, P_Δ93-126_-11S, or P_Δ39-57, Δ93-126_-11S and M-114, all resulted in luminescence comparable to transfection when P-11S was used. In contrast, when M-114 was transfected with P_ΔOD_-11S, luminescence signal was almost completely lost. These results indicate that, in transfected cells, P OD deletion, and as a consequence abrogated tetramerization, resulted in loss of interaction with M, whereas deletion of amino acids 39-57, 93-126 or both regions did not. Of note, analysis of the lysates of transfected 293T cells by Western blotting using anti-P or anti-M antibody confirmed correct protein expression with the fused NanoLUC fragments (Fig. 6E).

### Functional implications of P regions displaying direct M-binding

Next, we asked whether the identified M-binding domains on P were of functional significance. We analysed virus-like filament formation using full-length P or deletion constructs based on NMR results and on previous publications. Specifically, we checked full-length P, P_1-161_, P_ΔOD_, P_Δ39-57_, P_Δ93-110_ (deleted of M-binding site I), P_Δ113-126_ (deleted of M-binding site II), and P_Δ93-126_ (deleted of both sites I and II as identified by NMR) (Fig. 7A). BEAS-2B cells were transfected to express M, N, F and P_WT_ or various P deletion constructs, and the formation of RSV virus-like filaments was determined by confocal imaging and immuno-fluorescence after staining with anti-M and anti-P primary antibodies (Fig. 7B). Transfecting M, N, F and P resulted in pseudo-IBs and filament formation, and co-localization of M and P proteins in both (positive control, Fig. 7B, upper left). Transfecting M, N, F and P_ΔOD_ did not produce any virus-like filaments or pseudo-IBs, and M and P did not co-localize (Fig. 7B, lower left). Cells expressing P_Δ39-57_ failed to produce virus-like filaments, and although P was found in pseudo-IBs, M was not recruited to IBs (Fig. 7B, upper right). Transfection of P_Δ93-126_ resulted in virus-like filament formation (Fig. 7B, lower right), and both M and P proteins were found in virus-like filaments, similar to P_WT_, showing that sites I and II are not critical for these processes.

**Fig. 7:**
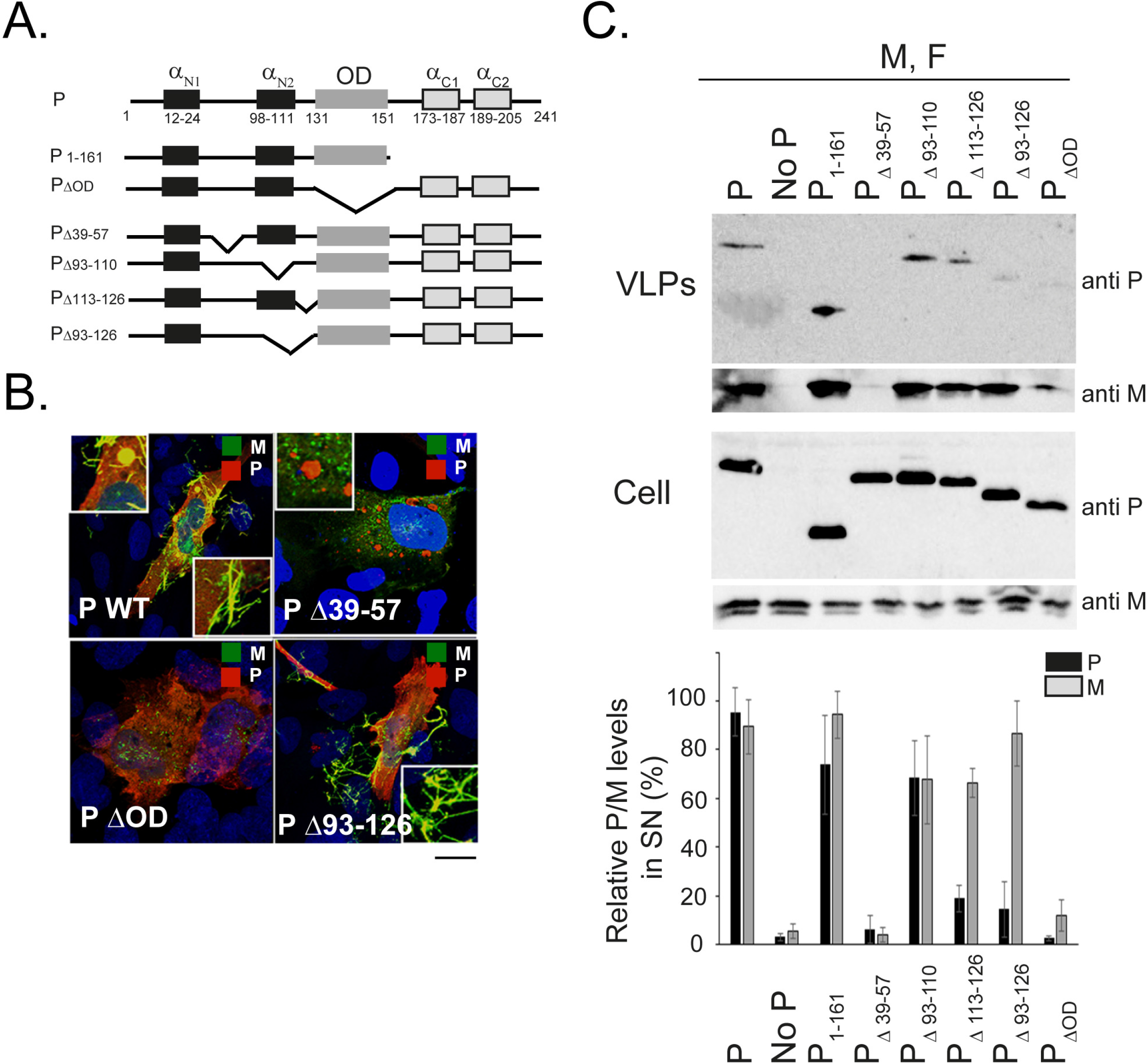
Functional analysis of P regions displaying direct M-binding. **A.** Scheme of the P protein secondary structure and the P constructs used. **B.** BEAS-2B cells were co-transfected with pcDNA3.1 plasmids expressing RSV M, N, F and P deletion mutants. Cells were fixed, permeabilized at 24 h post transfection, immunostained with anti-M and anti-P primary antibodies followed by Alexa Fluor secondary antibodies, and were analysed by confocal microscopy. Scale bars represent 10 μm. Merged images are zoomed in 3X. **C.** HEp-2 cells were co-transfected with pcDNA3.1 plasmids expressing RSV M, F and P_WT_ (lane 1, positive control) or pcDNA3.1 plasmids carrying RSV M, F and an empty pcDNA3.1 vector (lane 2, negative control) or with the indicated RSV P mutant constructs (lanes 3 to 8). At 48 h post transfection, VLPs (top) were isolated from the supernatant by pelleting of the clean supernatant through a sucrose cushion. Cell lysates (bottom) were generated using RIPA buffer. VLPs and cell lysates were then subjected to Western analysis using anti-P or anti-M polyclonal antibody. The amount of M and P protein in VLPs was quantified using the ImageJ software, and is presented as percentage (%) of M and P protein released when P_WT_ was used (100%). The graph is representative of four independent experiments. Data represents the % means and error bars represent standard deviation across 4 independent biological replicates.

Finally, we asked whether P deletion mutants could negatively affect VLP release. HEp-2 cells were transfected to express M, F, and various P constructs. The N protein was not included in the assay, since it is not required for VLP production (Fig. 1 and (17)). Cell lysates (soluble fractions) and the VLPs released into the media were analysed by Western blotting using anti-P and anti-M antibodies (Fig. 7C, upper panel). All P mutants were correctly expressed and were well detected by the anti-P antibody, as shown by the bands in the cell lysates. Quantification of relative M and P levels in the VLPs, revealed by WB, is shown in Fig. 7C, lower panel. When P_WT_ was expressed, VLP release was validated using anti-M antibodies, and P was detected in VLPs using anti-P antibodies (positive control). The absence of P prevented VLP release (negative control). The P_1-161_ construct resulted in VLP release comparable to P_WT_. Expression of P_ΔOD_ greatly reduced VLP release, as almost no M could be detected. P_Δ39-57_ abolished VLP release, as no M and P were detected. This is in agreement with our virus-like filament formation assay (Fig. 7B), where neither P_ΔOD_ nor P_Δ39-57_ induced filament formation. Transfection of P_Δ93-110_, P_Δ113-126_ or P_Δ93-126_ mutants did not abolish VLP release, as M was detected in the VLP fraction at comparable levels to P_WT_ with P_Δ93-126_, or slightly reduced with P_Δ93-110_ and P_Δ113-126_. P_Δ93-110_ incorporation into released VLPs was comparable to that of P_1-161_, i.e. with a 25-30% reduction relative to P_WT_. In contrast, transfection of P_Δ113-126_ and even more of P_Δ93-126_ greatly reduced detection of these mutants in released VLPs. This suggests that P_Δ113-126_ and P_Δ93-126_ mutants still supported VLP formation, in contrast to P_ΔOD_ and P_Δ39-57_, but could be more easily displaced form VLPs than P_Δ93-110_, P_1-161_ or P_WT_. This effect appears to be linked to the deletion of site II. It was not evidenced in the virus-like filament formation assay carried out with P_Δ93-126_ (Fig. 7D), most likely because of milder conditions.

## Discussion

### RSV M makes a direct interaction with RSV P

It was reported earlier that the three proteins, RSV P, M and F were sufficient for virus-like filament formation and VLP release (17). Our results confirm this minimal requirement (Fig. 1). A question that remained open was which interactions M needed to be involved in to promote assembly and budding of viral particles. For *Paramyxoviridae* it was shown that M proteins organize viral assembly by bridging between the glycoproteins and the RNPs, and that specific M-N interactions were required for the RNP to be packed into viral pseudo-particles (48). The interaction between the RNP and M proteins occurs most probably first in IBs and/or assembly granules (16), where M recruits the RNP, before they are transported to viral filaments formed on the cellular membrane.

Here we showed that M co-localized with P in pseudo-IBs, which are formed when N and P are present, as well as in virus-like filaments, even in the absence of N (Fig. 1). Our NanoLUC results showed that M could interact with P in the absence of other viral proteins in cells (Fig. 2). Gel shift assays performed with purified recombinant M and P fragments (Fig. 3C) confirmed M and P could directly interact. Moreover, our data indicated that the P domain responsible for this M interaction was located in fragment P_1-163_, comprising the N terminal domain and the OD of P (Fig. 3C and Fig. 6C). When M was co-expressed with P_1-161_, no pseudo-IBs were formed, as expected, since P_1-161_ lacks the C-terminus of P needed for interaction with RNA-N complexes and formation of IBs (11, 37, 45). However, virus-like filaments were formed, where M and P_1-161_ co-localized. This argues against the requirement of preliminary IB formation before virus-like filament assembly, as these were observed in cells transfected with RSV M, P, and F only. Furthermore, an interaction between RSV M and P strongly suggests that for RSV assembly, bridging between M and the RNP might be mediated by P.

### P displays multiple direct contact sites for M

According to our NMR data (Fig. 4 and 5), M binding to P could be achieved through multiple contact sites that are located within three regions, α_N2_ region (site I), the 115-125 region (site II) and the OD (Fig. 5). It must be noted that NMR experiments were carried out with the M-Y229A mutant, which stays dimeric in solution. The interactions observed by NMR therefore represent a P-M complex formed with dimeric M. The NMR results were corroborated *in vitro* by native gel interaction assays using both M WT and M-Y229A and N-terminally truncated P fragments (Fig. 3 and 6). Interestingly, neither sites I and II together but without the OD (P_1–126_ as compared to P_80–163_ or P_1–163_) nor the OD without sites I and II (P_120–160_ as compared to P_80–163_) seemed to be sufficient to bind M (Fig. 3C, 4, 5 and 6C). The split luciferase assay shed another light on the complexity of M-P interactions. The deletion of the internal N-terminal region spanning site I and II did not impair M binding *in cellula*, in contrast to the internal deletion of the OD, which completely impaired the M-P interaction under the same conditions (Fig. 6D). This could indicate that other short intrinsically disordered N-terminal motifs, similar in sequence to sites I or II, could complement the M-binding site, as they are closer to the OD than in P_WT_ due the 30-residue long truncation, or that cellular proteins help stabilize the M-P complex. Interestingly, as illustrated by the P_Δ39-57_ deletion mutant, highlighting the essential role of the 39-57 region (42), the ability to form an M-P complex *in cellula* is not correlated with virus-like filament formation or P-M co-localization on these virus-like filaments (Fig. 6D and 7B).

Although P OD is not sufficient to bind M, it seems to play a major role, possibly because of the higher level of structural organization provided by tetramerization. We previously showed that transient intra-protomer interactions take place in P (35), and the cryo-EM structures of *Pneumoviridae* L-P complexes show that the four protomers can engage each in different interaction in a single complex (32, 38, 39). This suggests that the P-M interactions are complex and probably cooperative, and that M could recognize one or several sites only formed when four P protomers are present.

As can be seen from *Pneumoviridae* P protein sequence alignment (Fig. 8), the N-terminal P region contains conserved motifs, in particular the N^0^-binding motif, the RVxF-like motif, which is the binding site for cellular PP1 phosphatase (41), and α_N2_, which is the binding site for M2-1 protein. This region also contains conserved clusters of negatively charged residues, notably just upstream and inside binding site II. As indicated by reduced M-P binding in high salt buffer, these residues could be important for interaction with M, which displays a large positively charged surface patch (29). Finally, the differences observed between *in vitro* and *in cellula* experiments could also depend on post-translational modifications of the proteins.

**Fig. 8:**
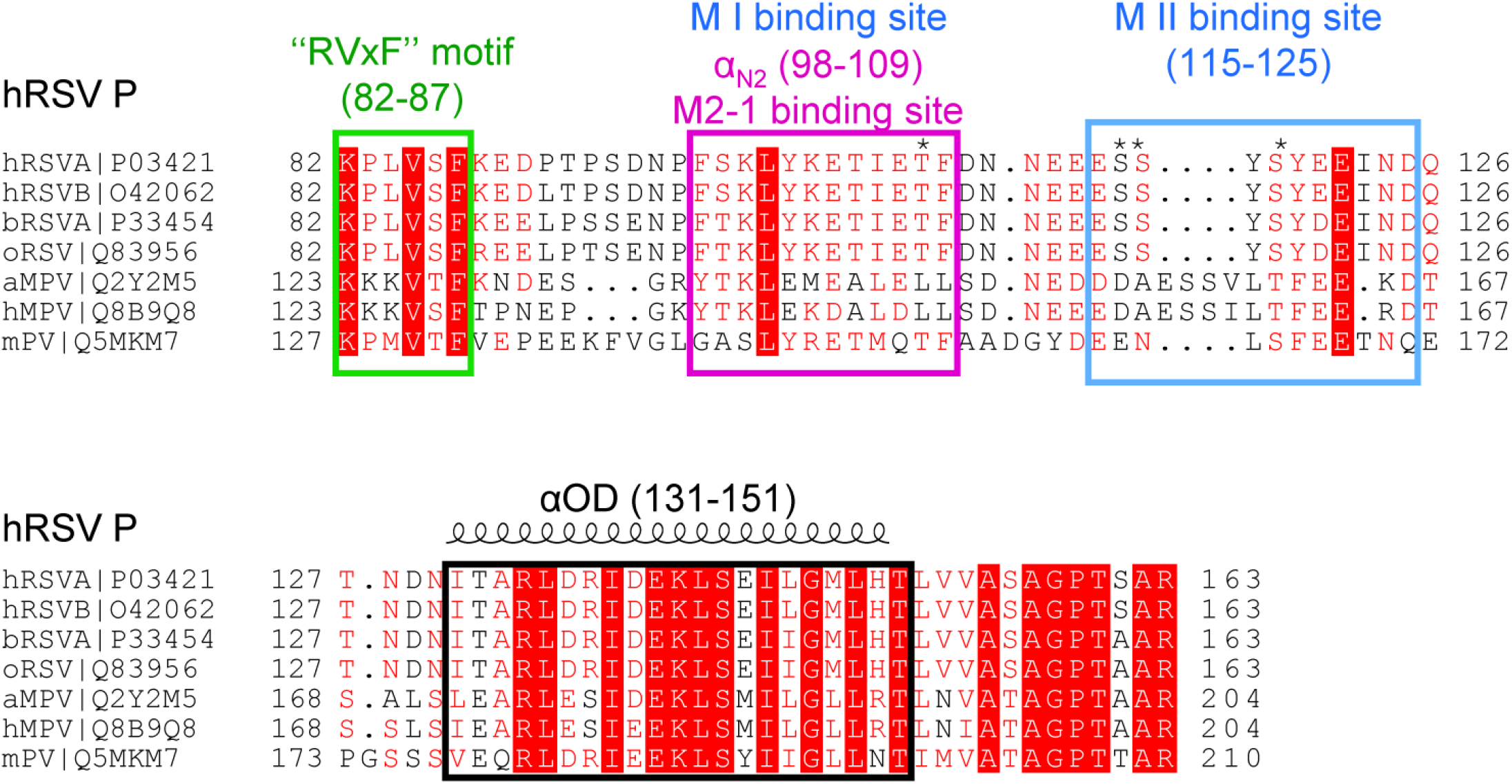
Sequence alignment of the central part of *Pneumoviridae* phosphoproteins. Sequence alignment for human (strains A and B), bovine and ovine respiratory syncytial virus, murine pneumonia virus, human and avian metapneumovirus P proteins was generated using T-coffee suite of program (58). Uniprot accession numbers are indicated next to each sequence and residue numbers are given for each sequence. Complete conservation is indicated with a white font on red background, and relative conservation is indicated by a red font. Conserved motifs and structure elements are annotated for hRSV P and boxed throughout the aligned sequences. The boundaries of the OD are from the cryo-EM structures of L-P complexes of hRSV and hMPV (32, 59). The “RVxF”-like motif that binds PP1 and the α_N2_ M2-1 binding site (41) are boxed in green and magenta, respectively. M binding sites I and M binding site II, as identified by NMR, are indicated or boxed in blue, respectively. Identified phosphorylation sites in the hRSV P (43, 54) are indicated by a star above the sequence.

### Functional relevance of direct binding between M and P

Our virus-like filament formation and VLP assays shed light on the functional implications of these interactions. Whereas the OD, and likely tetrameric organization provided by the OD, was clearly needed for VLP formation, deletion of sites I or II individually, or both at the same time, did not prevent virus-like filament production (Fig. 7B). However, reduced levels of P were detected in released VLPs, when both sites I and II were simultaneously deleted (Fig. 7C). This could be attributed to detachment of mutated P from released VLPs rather than defective recruitment to virus-like filaments, since co-localization was observed in these structures (Fig. 7B). This could also rise the question about the integrity of these VLPs.

M binding sites I and II thus rather appear to modulate P binding to M and to affect the final organization of the VLPs. The newly identified sites I and II also stand in contrast to the previously reported functional importance of the P region spanning residues 39-57 (42). In agreement with previously published data, we confirm that P_Δ39-57_ completely prevented filament formation and VLP budding (Fig. 7B and 7C). However, no perturbations were detected in the NMR signals in the 39-57 region of P in presence of M-Y229A (Fig. 4). The P-M interaction was still detected when residues 39-57 were deleted in transfected cells (Fig. 6D), indicating that this is probably not a direct binding region for M, at least not in its dimeric form. Dephosphorylation of the Ser/Thr-rich 39-57 region of P was reported to be required for VLP formation (42). This was the case for recombinant P protein produced in *E. coli*. As shown by our IF confocal microscopy experiments (Fig. 7B), M failed to co-localize to pseudo-IBs when P_Δ39-57_ was expressed. This could indicate that a third, possibly cellular factor is required for M recruitment to IBs. Some early reports have shown that M localization to IBs was mediated by an interaction with M2-1 (26, 27), and M interaction with M2-1 in cells was shown by co-IP (49). Cryo-EM analysis of RSV filamentous particles showed that M2-1 is located between M layer and RNP (14, 50, 51). However, our work here shows that M2-1 is not required for M localization into pseudo-IBs (Fig. 1). Moreover M and M2-1 did not interact in our NanoLUC assay (Fig. 2).

In conclusion, our virus-like filament formation and VLP assays show a strong functional relevance of P region 39-57, which is essential for M localization to IBs and budding process, but most probably not due to a direct P-M binding. In addition, we show the functional importance of the tetramerisation of P, occurring via the short OD region. The OD is probably part of the M interaction site with P, but must be complemented by the N-terminal region of P to bind M (Fig. 7).

### Possible role for RSV P as a switch between transcription and budding

Although the functional relevance of binding site I (α_N2_ region) could not be fully assessed in the framework of this study, it is striking that it completely overlaps with the binding site of transcription anti-termination factor M2-1, which is an essential factor for efficient RSV transcription (52) (Fig. 8). It is likely that RNA synthesis processes are frozen just before viral budding. However, such mechanism has not been clearly investigated to our knowledge. It is established that P plays a critical role in viral transcription, viral RNA synthesis and budding. In particular, interaction of P with M2-1 is critical for M2-1 recruitment to IBs (53). The overlap of binding sites for two essential proteins driving transcription (M2-1) on the one hand and virus assembly (M) on the other hand, could be part of a switch mechanism, where both proteins compete for this site. Interestingly, binding sites I and II contain four reported phosphorylation sites on T108 (54), S116, S117 and S119 (43), which are conserved among RSV P proteins (Fig. 8). Previously published data showed that P phosphorylation on residue T108 abolished P-M2-1 interaction (54). Phosphomimetic mutants of P serines 116, 117 and 119 (inside binding site II) were reported to significantly prevent virus budding (43). Moreover, P lacking residues 110-120 reduced budding, which was completely restored when these residues were added to P (42). P is known to be highly phosphorylated in purified virions, and the timing of appearance of phosphorylated P was shown to correspond to the release of RSV virions (55). We therefore suggest that the phosphorylation state of P sites I and II could regulate the switch between RSV transcription/replication and assembly.

In summary, our work further confirms that RSV P protein is a multifunctional protein playing different roles depending on its interactions with other viral proteins. We confirm here that P plays a key role in RSV assembly. We also bring evidence for a direct interaction between M and P, P OD being required for M-P interaction and sites I and II modulating P binding to M in VLPs by mechanisms yet to uncover.

## MATERIALS AND METHODS

### Plasmid constructs

pcDNA3.1 codon-optimized plasmids for mammalian expression encoding the RSV A2 M, P, N, F, and M2-1 proteins were a gift from Marty Moore, Emory University (56). Commercially made pciNanoLUC 114 and 11S vectors (GeneCust) were used to clone the RSV A2 codon-optimized M, P, N, and M2-1 constructs using standard PCR, digestion and ligation techniques. pcDNA P_1-126_ and P_1-161_ deletion mutants of P were obtained by introducing stop codons at the appropriate site in the coding sequence. All pcDNA3.1 P deletion mutants in the full-length construct were generated by using the Q5 site-directed mutagenesis kit (New England BioLabs), following the manufacturer recommendations. pciNanoLUC 11S P_Δ39-57_, P_Δ93–126_ P_Δ39–57, Δ93–126_ and P_ΔOD_ mutants were obtained by re-cloning the relevant constructs from pcDNA vector using standard PCR, digestion and ligation techniques. For expression and purification of recombinant P proteins, the previously described pGEX-P, and pGEX-P_1-126_, pGEX-P_1-163_, pGEX-P_ΔOD_, pGEX-P_127-241_ and pGEX-P_161-241_ plasmids were used (36). pGEX-P_OD_, pGEX-P_40-163_, pGEX-P_80-163_, pGEX-P_113-163_, and pGEX-P_120-160_ were generated using standard PCR, digestion and ligation techniques and introducing codons at the appropriate site in the coding sequence. It is important to note that pcDNA P_1-161_ and pGEX P_1-163_ were used for cellular expression or for bacterial expression respectively. For expression and purification of recombinant M protein, the previously described pCDF-M and pCDF-M_Y229A_ were used (28).

### Cell culture

HEp-2 (ATCC CCL-23™) and 293T cells were maintained in Dulbecco modified Eagle medium (eurobio) supplemented with 10% fetal calf serum (FCS; eurobio), 1% L-glutamine, and 1 % penicillin streptomycin. The transformed human bronchial epithelial cell line (BEAS-2B) (ATCC CRL-9609) was maintained in RPMI 1640 medium (eurobio) supplemented with 10% fetal calf serum (FCS; eurobio), 1% L-glutamine, and 1% penicillin-streptomycin. The cells were grown at 37°C in 5% CO2.

### Bacteria expression and purification of recombinant proteins

For M expression (WT and Y229A mutant), *E. coli* Rosetta 2 bacteria transformed with the pCDF-M plasmid were grown from fresh starter cultures in Luria-Bertani (LB) broth for 5 h at 32°C, followed by induction with 0.4 mM isopropylthi-galactoside (IPTG) for 4 h at 25°C. Cells were lysed by sonication (4 times for 20 s each time) and lysozyme (1 mg/ml; Sigma) in 50 mM NaH_2_PO_4_-Na_2_HPO_4_, 300 mM NaCl, pH 7.4, plus protease inhibitors (Roche), RNase (12 g/ml, Sigma), and 0.25% CHAPS {3-[(3-cholamidopropyl)-dimethylammonio]-1-propanesulfonate},. Lysates were clarified by centrifugation (23,425 g, 30 min, 4°C), and the soluble His_6_-M protein was purified on an Nickel sepharose column (HiTrap™ 5 ml IMAC HP; GE Healthcare). The bound protein was washed extensively with loading buffer plus 25 mM imidazole and eluted with a 25 to 250 mM imidazole gradient. M was concentrated to 2 ml using Vivaspin20 columns (SartoriusStedimBiotec) and purified on a HiLoad 10/600 Superdex S200 column (GE Healthcare) in 50 mM NaH_2_PO_4_-Na_2_HPO_4_, 300 mM NaCl, pH 7.4. The M peak was concentrated to 3 mg/ml using Vivaspin4 columns. The quality of protein samples was assessed by SDS-PAGE. Protein concentration was determined by measuring absorbance at 280 nm. For NMR interaction experiments a fresh preparation of M was dialyzed into NMR buffer.

For P expression, *E.coli* BL21 (DE3) bacteria transformed with pGEX-P derived plasmids were grown at 37°C for 8 h in LB. Protein expression was induced by adding one volume of fresh LB medium, 0.4 mM IPTG for 16 h at 28°C. Bacterial pellets were resuspended in lysis buffer (20 mM Tris/HCl pH 7.4, 60 mM NaCl, 1 mM EDTA, 1 mg/mL lysozyme, 1 mM DTT, 0,1% Triton X-100) supplemented with complete protease inhibitor cocktail (Roche) for 1 h on ice. Benzonase (Millipore) was then added and the lysate was incubated for 1 h at RT under rotation. The lysates were centrifuged at 4°C for 30 min at 10,000g. Glutathione-Sepharose 4B beads (GE Healthcare) were added to the clarified supernatants and the mixtures were incubated overnight at 4°C under rotation. The beads were washed with lysis buffer, three times with 1X phosphate-buffered saline (PBS) and then stored at 4°C in an equal volume of PBS.

For ^15^N-labeled P expression, bacteria were grown in minimal medium supplemented with ^15^NH_4_Cl (Eurisotop) as a ^15^N source. The GST tag was removed by thrombin (Millipore) cleavage and the cleaved product exchanged into NMR buffer (50 mM Na phosphate at pH 6.7, 150 mM NaCl). The quality of protein samples was assessed by SDS-PAGE. Protein concentration was determined by Bradford assay (Biorad) and checked by measuring absorbance at 280 nm for fragments containing tyrosine residues. It is given as protomer concentration in the case of P tetramers.

### NanoLUC interaction assay

Constructs expressing the NanoLUC subunits 114S and 11S were used (44). 293T cells were seeded at a concentration of 3×10^4^ cells per well in 48-well plate. After 24 h, cells were co-transfected in triplicate with 0.4 μg of total DNA (0.2 μg of each plasmid) using Lipofectamine 2000 (Invitrogen). 24 h post transfection cells were washed with PBS, and lysed for 1 h in room temperature using 50 μl NanoLUC lysis buffer (Promega). NanoLUC enzymatic activity was measured using the NanoLUC substrate (Promega). For each pair of plasmids, three normalized luminescence ratios (NLRs) were calculated as follows: the luminescence activity measured in cells transfected with the two plasmids (each viral protein fused to a different NanoLUC subunit) was divided by the sum of the luminescence activities measured in both control samples (each NanoLUC fused viral protein transfected with an plasmid expressing only the NanoLUC subunit) Data represents the mean ±SD of 4 independent experiments, each done in triplicate. Luminescence was measured using Infinite 200 Pro (Tecan, Männedorf, Switzerland).

### NMR spectroscopy

For NMR experiments, P and M solutions were mixed to obtain the desired molar ratio and concentrated to reach a concentration of 100 μM for M (WT and Y229A mutant).

NMR measurements were carried out in a Bruker Avance III spectrometer at a magnetic field of 18.8 T (800 MHz ^1^H frequency) equipped with a cryogenic TCI probe. The magnetic field was locked with 7 % ^2^H_2_O. The temperature was set to 288 K or 298 K. ^1^H-^15^N correlation spectra were acquired with a BEST-TROSY version. Spectra were processed with Topspin 4.0 (Bruker Biospin) and analyzed with CCPNMR 2.4 (57) software.

^1^H and ^15^N amide chemical shift assignment of full-length P and P_1-126_ and P_1-163_ fragments was done previously (46). Amide chemical shift assignment of ^15^N-labeled P_113-160_ and of ^15^N-labeled P_120-160_ was done by recording a ^15^N-separated NOESY-HSQC spectrum with 80 ms mixing time. Sequential information was retrieved through H_Ni_-H_Ni-1_,H_Ni_-H_Ni+1_ and H_Ni_-H_αi-1_ correlations.

### Virus-like filament/particle formation

Over night cultures of BEAS-2B cells seeded at 4 10^5^ cells/well in 6-well plates (on a 16-mm micro-cover glass for immunostaining) were transfected with pcDNA3.1 codon-optimized plasmids (0.4 μg each) carrying the RSV A2 WT or deletion/mutant P protein along with pcDNA3.1 codon-optimized plasmids carrying RSV A2 M, N, and F using Lipofectamine 2000 (Invitrogen) according to the manufacturer’s recommendations. Cells were fixed at 24 h post transfection, immunostained, and imaged as described below. For VLP formation, over-night cultures of HEp-2 cells seeded at 4 10^5^ cells/well in 6-well plates were transfected as described above. Released VLPs were harvested from the supernatant; the supernatant was clarified of cell debris by centrifugation (1,300 g, 10 min, 4°C) and pelleted through a 20% sucrose cushion (13,500 g, 90 min, 4°C). Cells were lysed in radio immune precipitation assay (RIPA) buffer. Cellular lysates and VLP pellets were dissolved in Laemmli buffer and subjected to Western analysis.

### Immunostaining and imaging

Cells were fixed with 4% paraformaldehyde in PBS for 10 min, blocked with 3% BSA in 0.2% Triton X-100–PBS for 10 min, and immunostained with monoclonal anti-M (1:200; a gift from Mariethe Ehnlund, Karolinska Institute, Sweden), polyclonal anti-N (1:5000; (30)), polyclonal anti-P (1:500; (30)) or monoclonal anti-P (1:100; a gift from Jose A. Melero, Madrid, Spain) antibodies, followed by species-specific secondary antibodies conjugated to Alexa Fluor 488 and Alexa Fluor 568 (1: 1,000; Invitrogen). Images were obtained using the White Light laser SP8 (Leica Microsystems, Wetzlar, Germany) confocal microscope at a nominal magnification of 63. Images were acquired using the Leica Application Suite X (LAS X) software.

### SDS-PAGE and Western analysis

Protein samples were separated by electrophoresis on 12% polyacrylamide gels in Tris-glycine buffer. All samples were boiled for 3 min prior to electrophoresis. Proteins were then transferred to a nitrocellulose membrane (RocheDiagnostics). The blots were blocked with 5% non fat milk in Tris-buffered saline (pH 7.4), followed by incubation with rabbit anti-P antiserum (1:5,000) (30), rabbit anti N antiserum (1:5,000) (30), rabbit anti M antiserum (1:1,000), or rabbit anti M2-1 antiserum (1:2,000) (30), and horseradish peroxidase (HRP)-conjugated donkey anti-rabbit (1:5,000) antibodies (P.A.R.I.S.). Western blots were developed using freshly prepared chemiluminescent substrate (100 mM TrisHCl, pH 8.8, 1.25 mM luminol, 0.2 mM p-coumaric acid, 0.05% H_2_O_2_) and exposed using BIO-RAD ChemiDoc™ Touch Imaging System.

### Generation of M antiserum

Polyclonal anti M serum was prepared by immunizing a rabbit three times at 2 week intervals using purified His-fusion proteins (100 mg) for each immunization. The first and second immunizations were administered subcutaneously in 1 ml Freund’s complete and Freund’s incomplete adjuvant (Difco), respectively. The third immunization was done intramuscularly in Freund’s incomplete adjuvant. Animals were bled 10 days after the third immunization.

### Native gel

Protein samples were separated by electrophoresis on 1% agarose gel in TBE (Tris Borate EDTA) buffer. 50 μg M and x 10 μg P were incubated in PBS buffer (pH 7.4) for 20 min in RT. Samples were mixed with 50% sucrose and run for 2 h at 80 V, following by staining with Amido Black.

## Acknowledgements

We thank Benoit Maury and CYMAGES Imaging facility (Département biotechnologie Santé, Université Versailles Saint Quentin, France) for their support & assistance with the confocal imaging. We thank Damien Vitour (E’quipe du laboratoire d’immunologie de Seppic and the animal facilities, ANSES Maisons-Alfort) for rabbit immunization with recombinant M. This work was carried out with the financial support of the French Agence Nationale de la Recherche DecRisP (ANR-19_CE11_0017).

## References

1. Group TPERfCHPS. 2019. Causes of severe pneumonia requiring hospital admission in children without HIV infection from Africa and Asia: the PERCH multi-country case-control study. Lancet 394:757–79.

2. Grimprel E. 2001. [Epidemiology of infant bronchiolitis in France]. Arch Pediatr 8 Suppl 1:83S–92S.

3. Olszewska W, Openshaw P. 2009. Emerging drugs for respiratory syncytial virus infection. Expert Opin Emerg Drugs 14:207–17.

4. Mazur NI, Higgins D, Nunes MC, Melero JA, Langedijk AC, Horsley N, Buchholz UJ, Openshaw PJ, McLellan JS, Englund JA, Mejias A, Karron RA, Simoes EA, Knezevic I, Ramilo O, Piedra PA, Chu HY, Falsey AR, Nair H, Kragten-Tabatabaie L, Greenough A, Baraldi E, Papadopoulos NG, Vekemans J, Polack FP, Powell M, Satav A, Walsh EE, Stein RT, Graham BS, Bont LJ. 2018. The respiratory syncytial virus vaccine landscape: lessons from the graveyard and promising candidates. Lancet Infect Dis 18:e295–e311.

5. Schmidt MR, McGinnes-Cullen LW, Kenward SA, Willems KN, Woodland RT, Morrison TG. 2014. Modification of the respiratory syncytial virus f protein in virus-like particles impacts generation of B cell memory. J Virol 88:10165–76.

6. Lee YN, Hwang HS, Kim MC, Lee YT, Lee JS, Moore ML, Kang SM. 2015. Recombinant influenza virus expressing a fusion protein neutralizing epitope of respiratory syncytial virus (RSV) confers protection without vaccine-enhanced RSV disease. Antiviral Res 115:1–8.

7. Bachi T, Howe C. 1973. Morphogenesis and ultrastructure of respiratory syncytial virus. J Virol 12:1173–80.

8. Afonso CL, Amarasinghe GK, Banyai K, Bao Y, Basler CF, Bavari S, Bejerman N, Blasdell KR, Briand FX, Briese T, Bukreyev A, Calisher CH, Chandran K, Cheng J, Clawson AN, Collins PL, Dietzgen RG, Dolnik O, Domier LL, Durrwald R, Dye JM, Easton AJ, Ebihara H, Farkas SL, Freitas-Astua J, Formenty P, Fouchier RA, Fu Y, Ghedin E, Goodin MM, Hewson R, Horie M, Hyndman TH, Jiang D, Kitajima EW, Kobinger GP, Kondo H, Kurath G, Lamb RA, Lenardon S, Leroy EM, Li CX, Lin XD, Liu L, Longdon B, Marton S, Maisner A, Muhlberger E, Netesov SV, Nowotny N, et al. 2016. Taxonomy of the order Mononegavirales: update 2016. Arch Virol 161:2351–60.

9. Rincheval V, Lelek M, Gault E, Bouillier C, Sitterlin D, Blouquit-Laye S, Galloux M, Zimmer C, Eleouet JF, Rameix-Welti MA. 2017. Functional organization of cytoplasmic inclusion bodies in cells infected by respiratory syncytial virus. Nat Commun 8:563.

10. Garcia J, Garcia-Barreno B, Vivo A, Melero JA. 1993. Cytoplasmic inclusions of respiratory syncytial virus-infected cells: formation of inclusion bodies in transfected cells that coexpress the nucleoprotein, the phosphoprotein, and the 22K protein. Virology 195:243–7.

11. Galloux M, Risso-Ballester J, Richard CA, Fix J, Rameix-Welti MA, Eleouet JF. 2020. Minimal Elements Required for the Formation of Respiratory Syncytial Virus Cytoplasmic Inclusion Bodies In Vivo and In Vitro. MBio 11.

12. Roberts SR, Compans RW, Wertz GW. 1995. Respiratory syncytial virus matures at the apical surfaces of polarized epithelial cells. J Virol 69:2667–73.

13. Bajorek M, Caly L, Tran KC, Maertens GN, Tripp RA, Bacharach E, Teng MN, Ghildyal R, Jans DA. 2014. The Thr205 phosphorylation site within respiratory syncytial virus matrix (M) protein modulates M oligomerization and virus production. J Virol 88:6380–93.

14. Ke Z, Dillard RS, Chirkova T, Leon F, Stobart CC, Hampton CM, Strauss JD, Rajan D, Rostad CA, Taylor JV, Yi H, Shah R, Jin M, Hartert TV, Peebles RS, Jr., Graham BS, Moore ML, Anderson LJ, Wright ER. 2018. The Morphology and Assembly of Respiratory Syncytial Virus Revealed by Cryo-Electron Tomography. Viruses 10.

15. Vanover D, Smith DV, Blanchard EL, Alonas E, Kirschman JL, Lifland AW, Zurla C, Santangelo PJ. 2017. RSV glycoprotein and genomic RNA dynamics reveal filament assembly prior to the plasma membrane. Nat Commun 8:667.

16. Blanchard EL, Braun MR, Lifland AW, Ludeke B, Noton SL, Vanover D, Zurla C, Fearns R, Santangelo PJ. 2020. Polymerase-tagged respiratory syncytial virus reveals a dynamic rearrangement of the ribonucleocapsid complex during infection. PLoS Pathog 16:e1008987.

17. Meshram CD, Baviskar PS, Ognibene CM, Oomens AG. 2016. The Respiratory Syncytial Virus Phosphoprotein, Matrix Protein, and Fusion Protein Carboxy-Terminal Domain Drive Efficient Filamentous Virus-Like Particle Formation. J Virol 90:10612–10628.

18. Shaikh FY, Cox RG, Lifland AW, Hotard AL, Williams JV, Moore ML, Santangelo PJ, Crowe JE, Jr. 2012. A critical phenylalanine residue in the respiratory syncytial virus fusion protein cytoplasmic tail mediates assembly of internal viral proteins into viral filaments and particles. MBio 3.

19. McLellan JS, Chen M, Leung S, Graepel KW, Du X, Yang Y, Zhou T, Baxa U, Yasuda E, Beaumont T, Kumar A, Modjarrad K, Zheng Z, Zhao M, Xia N, Kwong PD, Graham BS. 2013. Structure of RSV fusion glycoprotein trimer bound to a prefusion-specific neutralizing antibody. Science 340:1113–7.

20. McLellan JS, Yang Y, Graham BS, Kwong PD. 2011. Structure of respiratory syncytial virus fusion glycoprotein in the postfusion conformation reveals preservation of neutralizing epitopes. J Virol 85:7788–96.

21. Swanson KA, Settembre EC, Shaw CA, Dey AK, Rappuoli R, Mandl CW, Dormitzer PR, Carfi A. 2011. Structural basis for immunization with postfusion respiratory syncytial virus fusion F glycoprotein (RSV F) to elicit high neutralizing antibody titers. Proc Natl Acad Sci U S A 108:9619–24.

22. Ghildyal R, Ho A, Jans DA. 2006. Central role of the respiratory syncytial virus matrix protein in infection. FEMS Microbiol Rev 30:692–705.

23. Kiss G, Chen X, Brindley MA, Campbell P, Afonso CL, Ke Z, Holl JM, Guerrero-Ferreira RC, Byrd-Leotis LA, Steel J, Steinhauer DA, Plemper RK, Kelly DF, Spearman PW, Wright ER. 2014. Capturing enveloped viruses on affinity grids for downstream cryo-electron microscopy applications. Microsc Microanal 20:164–74.

24. Harrison MS, Sakaguchi T, Schmitt AP. 2010. Paramyxovirus assembly and budding: building particles that transmit infections. Int J Biochem Cell Biol 42:1416–29.

25. Mitra R, Baviskar P, Duncan-Decocq RR, Patel D, Oomens AG. 2012. The human respiratory syncytial virus matrix protein is required for maturation of viral filaments. J Virol 86:4432–43.

26. Ghildyal R, Mills J, Murray M, Vardaxis N, Meanger J. 2002. Respiratory syncytial virus matrix protein associates with nucleocapsids in infected cells. J Gen Virol 83:753–7.

27. Li D, Jans DA, Bardin PG, Meanger J, Mills J, Ghildyal R. 2008. Association of respiratory syncytial virus M protein with viral nucleocapsids is mediated by the M2-1 protein. J Virol 82:8863–70.

28. Forster A, Maertens GN, Farrell PJ, Bajorek M. 2015. Dimerization of matrix protein is required for budding of respiratory syncytial virus. J Virol 89:4624–35.

29. Money VA, McPhee HK, Mosely JA, Sanderson JM, Yeo RP. 2009. Surface features of a Mononegavirales matrix protein indicate sites of membrane interaction. Proc Natl Acad Sci U S A 106:4441–6.

30. Castagne N, Barbier A, Bernard J, Rezaei H, Huet JC, Henry C, Da Costa B, Eleouet JF. 2004. Biochemical characterization of the respiratory syncytial virus P-P and P-N protein complexes and localization of the P protein oligomerization domain. J Gen Virol 85:1643–53.

31. Llorente MT, Taylor IA, Lopez-Vinas E, Gomez-Puertas P, Calder LJ, Garcia-Barreno B, Melero JA. 2008. Structural properties of the human respiratory syncytial virus P protein: evidence for an elongated homotetrameric molecule that is the smallest orthologue within the family of paramyxovirus polymerase cofactors. Proteins 72:946–58.

32. Gilman MSA, Liu C, Fung A, Behera I, Jordan P, Rigaux P, Ysebaert N, Tcherniuk S, Sourimant J, Eleouet JF, Sutto-Ortiz P, Decroly E, Roymans D, Jin Z, McLellan JS. 2019. Structure of the Respiratory Syncytial Virus Polymerase Complex. Cell 179:193–204 e14.

33. Simabuco FM, Asara JM, Guerrero MC, Libermann TA, Zerbini LF, Ventura AM. 2011. Structural analysis of human respiratory syncytial virus p protein: identification of intrinsically disordered domains. Braz J Microbiol 42:340–5.

34. Noval MG, Esperante SA, Molina IG, Chemes LB, Prat-Gay G. 2016. Intrinsic Disorder to Order Transitions in the Scaffold Phosphoprotein P from the Respiratory Syncytial Virus RNA Polymerase Complex. Biochemistry 55:1441–54.

35. Pereira N, Cardone C, Lassoued S, Galloux M, Fix J, Assrir N, Lescop E, Bontems F, Eleouet JF, Sizun C. 2017. New Insights into Structural Disorder in Human Respiratory Syncytial Virus Phosphoprotein and Implications for Binding of Protein Partners. J Biol Chem 292:2120–2131.

36. Galloux M, Gabiane G, Sourimant J, Richard CA, England P, Moudjou M, Aumont-Nicaise M, Fix J, Rameix-Welti MA, Eleouet JF. 2015. Identification and characterization of the binding site of the respiratory syncytial virus phosphoprotein to RNA-free nucleoprotein. J Virol 89:3484–96.

37. Tran TL, Castagne N, Bhella D, Varela PF, Bernard J, Chilmonczyk S, Berkenkamp S, Benhamo V, Grznarova K, Grosclaude J, Nespoulos C, Rey FA, Eleouet JF. 2007. The nine C-terminal amino acids of the respiratory syncytial virus protein P are necessary and sufficient for binding to ribonucleoprotein complexes in which six ribonucleotides are contacted per N protein protomer. J Gen Virol 88:196–206.

38. Sourimant J, Rameix-Welti MA, Gaillard AL, Chevret D, Galloux M, Gault E, Eleouet JF. 2015. Fine mapping and characterization of the L-polymerase-binding domain of the respiratory syncytial virus phosphoprotein. J Virol 89:4421–33.

39. Cao D, Gao Y, Roesler C, Rice S, D’Cunha P, Zhuang L, Slack J, Domke M, Antonova A, Romanelli S, Keating S, Forero G, Juneja P, Liang B. 2020. Cryo-EM structure of the respiratory syncytial virus RNA polymerase. Nat Commun 11:368.

40. Selvaraj M, Yegambaram K, Todd E, Richard CA, Dods RL, Pangratiou GM, Trinh CH, Moul SL, Murphy JC, Mankouri J, Eleouet JF, Barr JN, Edwards TA. 2018. The Structure of the Human Respiratory Syncytial Virus M2-1 Protein Bound to the Interaction Domain of the Phosphoprotein P Defines the Orientation of the Complex. MBio 9.

41. Richard CA, Rincheval V, Lassoued S, Fix J, Cardone C, Esneau C, Nekhai S, Galloux M, Rameix-Welti MA, Sizun C, Eleouet JF. 2018. RSV hijacks cellular protein phosphatase 1 to regulate M2-1 phosphorylation and viral transcription. PLoS Pathog 14:e1006920.

42. Meshram CD, Oomens AGP. 2019. Identification of a human respiratory syncytial virus phosphoprotein domain required for virus-like-particle formation. Virology 532:48–54.

43. Lu B, Ma CH, Brazas R, Jin H. 2002. The major phosphorylation sites of the respiratory syncytial virus phosphoprotein are dispensable for virus replication in vitro. J Virol 76:10776–84.

44. Dixon AS, Schwinn MK, Hall MP, Zimmerman K, Otto P, Lubben TH, Butler BL, Binkowski BF, Machleidt T, Kirkland TA, Wood MG, Eggers CT, Encell LP, Wood KV. 2016. NanoLuc Complementation Reporter Optimized for Accurate Measurement of Protein Interactions in Cells. ACS Chem Biol 11:400–8.

45. Galloux M, Tarus B, Blazevic I, Fix J, Duquerroy S, Eleouet JF. 2012. Characterization of a viral phosphoprotein binding site on the surface of the respiratory syncytial nucleoprotein. J Virol 86:8375–87.

46. Pereira N, Cardone C, Lassoued S, Galloux M, Fix J, Assrir N, Lescop E, Bontems F, Eleouet JF, Sizun C. 2017. New Insights into Structural Disorder in Human Respiratory Syncytial Virus Phosphoprotein and Implications for Binding of Protein Partners. The Journal of biological chemistry 292:2120–2131.

47. Richard CA, Rincheval V, Lassoued S, Fix J, Cardone C, Esneau C, Nekhai S, Galloux M, Rameix-Welti MA, Sizun C, Eleouet JF. 2018. RSV hijacks cellular protein phosphatase 1 to regulate M2-1 phosphorylation and viral transcription. PLoS Pathogens 14:e1006920.

48. Ray G, Schmitt PT, Schmitt AP. 2016. C-Terminal DxD-Containing Sequences within Paramyxovirus Nucleocapsid Proteins Determine Matrix Protein Compatibility and Can Direct Foreign Proteins into Budding Particles. J Virol 90:3650–60.

49. Kipper S, Hamad S, Caly L, Avrahami D, Bacharach E, Jans DA, Gerber D, Bajorek M. 2015. New host factors important for respiratory syncytial virus (RSV) replication revealed by a novel microfluidics screen for interactors of matrix (M) protein. Mol Cell Proteomics 14:532–43.

50. Kiss G, Holl JM, Williams GM, Alonas E, Vanover D, Lifland AW, Gudheti M, Guerrero-Ferreira RC, Nair V, Yi H, Graham BS, Santangelo PJ, Wright ER. 2014. Structural analysis of respiratory syncytial virus reveals the position of M2-1 between the matrix protein and the ribonucleoprotein complex. J Virol 88:7602–17.

51. Liljeroos L, Krzyzaniak MA, Helenius A, Butcher SJ. 2013. Architecture of respiratory syncytial virus revealed by electron cryotomography. Proc Natl Acad Sci U S A 110:11133–8.

52. Fearns R, Collins PL. 1999. Role of the M2-1 transcription antitermination protein of respiratory syncytial virus in sequential transcription. J Virol 73:5852–64.

53. Blondot ML, Dubosclard V, Fix J, Lassoued S, Aumont-Nicaise M, Bontems F, Eleouet JF, Sizun C. 2012. Structure and functional analysis of the RNA- and viral phosphoprotein-binding domain of respiratory syncytial virus M2-1 protein. PLoS Pathog 8:e1002734.

54. Asenjo A, Calvo E, Villanueva N. 2006. Phosphorylation of human respiratory syncytial virus P protein at threonine 108 controls its interaction with the M2-1 protein in the viral RNA polymerase complex. J Gen Virol 87:3637–42.

55. Lambert DM, Hambor J, Diebold M, Galinski B. 1988. Kinetics of synthesis and phosphorylation of respiratory syncytial virus polypeptides. J Gen Virol 69 (Pt 2):313–23.

56. Hotard AL, Shaikh FY, Lee S, Yan D, Teng MN, Plemper RK, Crowe JE, Jr., Moore ML. 2012. A stabilized respiratory syncytial virus reverse genetics system amenable to recombination-mediated mutagenesis. Virology 434:129–36.

57. Vranken WF, Boucher W, Stevens TJ, Fogh RH, Pajon A, Llinas M, Ulrich EL, Markley JL, Ionides J, Laue ED. 2005. The CCPN data model for NMR spectroscopy: development of a software pipeline. Proteins 59:687–96.

58. Notredame C, Higgins DG, Heringa J. 2000. T-Coffee: A novel method for fast and accurate multiple sequence alignment. J Mol Biol 302:205–17.

59. Pan J, Qian X, Lattmann S, El Sahili A, Yeo TH, Jia H, Cressey T, Ludeke B, Noton S, Kalocsay M, Fearns R, Lescar J. 2020. Structure of the human metapneumovirus polymerase phosphoprotein complex. Nature 577:275–279.

